# Phylogenomics and body shape morphometrics reveal recent diversification in the goatfishes (Syngnatharia: Mullidae)

**DOI:** 10.1101/2022.04.12.488079

**Authors:** Chloe M. Nash, Linnea L. Lungstrom, Lily C. Hughes, Mark W. Westneat

**Author notes:** **Corresponding Author:** Chloe Nash.

## Abstract

Clades of marine fishes exhibit many patterns of diversification, ranging from relatively constant throughout time to rapid changes in the rates of speciation and extinction. The goatfishes (Syngnatharia: Mullidae) are a family of marine, reef associated fishes with a relatively recent origin, distributed globally in tropical and temperate waters. Despite their abundance and economic importance, the goatfishes remain one of the few coral reef families for which the species level relationships have not been examined using genomic techniques. Here we use phylogenomic analysis of ultra-conserved elements (UCE) and exon data to resolve a well-supported, time-calibrated phylogeny for 72 species of goatfishes, supporting a recent crown age of the goatfishes at 21.9 million years ago. We used this framework to test hypotheses about the associations among body shape morphometrics, taxonomy, and phylogeny, as well as to explore relative diversification rates across the phylogeny. Body shape was strongly associated with generic-level taxonomy of goatfishes, with morphometric analyses showing evidence for high phylogenetic signal across all morphotypes. Rates of diversification in this clade reveal a recent sharp increase in lineage accumulation, with 92% of the goatfish species sampled across all clades and major body plans having originated in just the past 5 million years. We suggest that habitat diversity in the early Pliocene oceans and the generalist ecology of goatfishes are key factors in the unusual evolutionary tempo of the family Mullidae.

## 1. Introduction

Shifts in the tempo and mode of evolution across lineages are a major force in generating global biodiversity (Simpson 1945; Gould and Eldredge 1977). Modern approaches have yielded novel evolutionary insights in many regions of the Tree of Life through the detection of variability in patterns of species diversification across large phylogenetic scales (Pennell et al. 2012; Stadler 2013; Donoghue and Sanderson 2015; Graham et al. 2018; Rabosky et al. 2018). This variation in the timing and rates of speciation and extinction is often the result of habitat fragmentation and dispersal capabilities, which can alter the amount of gene flow between populations (Kisel et al. 2011). As the largest group of living vertebrates, fishes have been a useful system to explore the impact of variable rates of diversification on current patterns of diversity (i.e. Alfaro et al. 2009; McMahan et al. 2013; Rabosky et al. 2013; Santini et al. 2013; Melo et al. 2021).

Heightened diversification rates in temperate fishes have been linked to particular geologic or climatic events (Rabosky et al. 2018), and likewise, bursts of evolution in reef fishes are linked to temporal patterns of expansion of coral reefs (Leprieur et al. 2021). The tempo and mode of diversification in fishes varies widely across taxonomic groups. Rates of diversification may be elevated towards the crown, as in Caribbean hamlets (Hench et al. 2022); towards the root, as in the case of neotropical cichlids (López-Fernández et al. 2013); or a relatively constant rate through the phylogenetic history of a group, such as in the damselfishes (McCord et al. 2021). Our ability to infer variable rates of lineage accumulation depends on our confidence in the underlying phylogeny and estimation of divergence times. Fortunately, genome-wide datasets that can greatly improve the resolution of phylogenetic relationships are now attainable for non-model organisms.

The goatfishes (Syngnatharia: Mullidae) are a diverse, globally distributed family of fishes found in close association with coral reef ecosystems. First described by Linnaeus in *Systema Naturae* (1758), the family now contains 102 valid species in 6 genera (Fricke et al. 2022). There have been several genus-level phylogenetic studies of goatfishes that provide the necessary framework to examine the complex genetic and morphological relationships among the many closely related species (Turan 2006; Keskin and Can 2009; Uiblein and Heemstra 2010; Uiblein 2011; Uiblein and Gouws 2015). However, despite their common occurrence on reefs worldwide, the goatfishes lack a comprehensive species-level molecular phylogeny, unlike many other charismatic reef fish families.

Most coral reef fish families are estimated to have originated in the Eocene, with subsequent diversification having occurred through the past 50-60 million years (Cowman and Bellwood 2011; Bannikov 2014; Bellwood et al. 2017). Although prior research on higher-level phylogenetics of fishes have estimated the origin of goatfishes to have occurred during this time period (i.e. Betancur-R et al. 2013; Near et al. 2013; Rabosky et al. 2018), recent phylogenomic studies that included a sample of goatfishes inferred a much more recent root node age of ∼18 Ma for the Mullidae (Santaquiteria et al. 2021). This raises intriguing questions about the timing of goatfish origins and the tempo of diversification that require a well-sampled time-calibrated phylogeny of the family to answer.

Goatfish taxonomy has been primarily based on morphological characters, such as dentition, fin ray counts, and body shape (i.e. Uiblein and Heemstra 2011; Uiblein and McGrouther 2012; Uiblein and Gouws 2013; Bos 2014). These morphological traits have been used to broadly examine phylogenetic relationships among genera, however the resolution of the data resulted in several large polytomies (Kim 2002). The main synapomorphy that unites goatfishes is a pair of highly specialized hyoid barbels, which are fleshy extensions capable of chemoreception and prey excavation (Gosline 1985). Goatfishes use these barbels to detect and resuspend otherwise inaccessible food sources within the benthic substrate (Gosline 1985). This unique behavior has been shown to play an important role in maintaining coral reef diversity and community composition (Lukoschek and McCormick 2000), in addition to acting as indicators of coral reef health (Uiblein 2007; Russ et al. 2015). These traits may provide the necessary framework to examine the complex genetic and morphological relationships among many closely related species and reiterate the need for a comprehensive phylogeny to improve our understanding of goatfish evolution. The integration of phylogenomics with geometric morphometrics will allow us to examine morphological evolution associated with lineage diversification across the goatfishes.

The central aims of this study are to 1) infer the phylogenetic relationships among species, 2) test hypotheses about the congruence between body shape morphology, taxonomy, and phylogenetic placement, and 3) examine the tempo of phylogenetic and morphological diversification across species of goatfishes. To achieve the primary aim, we use a robust genomic dataset to infer the first molecular phylogeny of goatfishes using combination of ultra-conserved elements (UCEs), exons, and ribosomal sequences in order to include the most species possible from throughout the family. UCEs and exon capture have been shown to be effective in resolving evolutionary relationships at different degrees of divergence, although genome coverage inversely decreases with the amount of divergence among species (Faircloth et al. 2012; Bragg et al. 2016). Additionally, we examine the evolution of body shape using a comprehensive geometric morphometric dataset to assess the phylogenetic affinities of goatfish morphology.

## 2. Methods

### 2.1 Taxon Sampling

We developed a comprehensive data matrix of 72 species of Mullidae, representing all genera in the family. We include 38 published UCE assemblies for Mullidae from two recent studies (Longo et al. 2017; Santaquiteria et al. 2021), as well as the full genome of *Mullus surmuletus* (Fietz et al. 2020). We expanded this taxon sampling by sequencing an additional 22 species of Mullidae (Table S1) using both acanthomorph UCE (Alfaro et al. 2018) and sygnatharian-specific exon capture sequencing probes (Hughes et al. 2021). To further increase taxon sampling, we compiled a matrix of 7 mitochondrial loci (*12S*, *16S*, *ATP6*, *COI*, *COII*, *cytB*, and *nd2*) and one nuclear locus (*rhod*) to incorporate an additional 11 species from previously published datasets available on NCBI (Table S2). Sequences for an additional 14 species were additionally from Longo *et al*. as representative outgroup lineages within Syngnatharia. Species identification was confirmed for all samples through comparison of COI with sequences available in the BOLD database. One specimen previously identified as *Upeneus guttatus* was updated to *Upeneus willwhite* given the recent reclassification of the sampled specimen (Uiblein and Motomura 2021). Additionally, we updated the name of a specimen initially identified as *Upeneus taeniopterus* to *Upeneus suahelicus* based on taxonomic input by Franz Uiblein (Uiblein and Gouws 2015, Table S1). However, given its phylogenetic distinctivness from the other *U. suahelicus* included in this study, we kept its classification as a distinct taxonomic unit.

### 2.2 Library Preparation and Sequencing

Genomic DNA from 22 goatfish samples was extracted from preserved tissues in a 96-well plate format on a GenePrep (Autogen) at the Smithsonian Institution Laboratory of Analytic Biology (Washington, DC), following manufacturer’s instructions. Extraction quality was assessed via visual inspections of genomic DNA on a 1.5% agarose gel. Illumina library preparation and sequence capture of both UCEs (Acanthomorph 1Kv1; Alfaro et al. 2018) and single-copy exons (Hughes et al. 2021) were conducted at Arbor Biosciences (Ann Arbor, MI). Libraries were sequenced on one lane of an Illumina HiSeq 4000 at the University of Chicago Genomics Facility as 100-bp paired-end reads. The sequence reads generated here have been deposited on NCBI’s SRA database under BioProject PRJNA824087.

### 2.3 UCE Data Assembly and Bioinformatic Processing

We assembled the genomic data using the phyluce 1.6.8 pipeline (Faircloth 2016). Read data was trimmed and cleaned using the default settings for illumiprocessor 2.0.9 (Faircloth 2013), which uses trimmomatic 0.39-1 (Bolger et al. 2014), to remove adapter contamination and low-quality bases. After adaptor quality and trimming of the 75 raw sequences, an average of 21,641 contigs were assembled in Trinity 2.8.5 (Grabherr et al. 2011) and were matched to the UCE Acanthomorph probe set (Faircloth et al. 2013). We recovered a total of 1,188 UCE loci with an average of 543 loci per sample. Loci were concatenated using mafft v7 (Katoh et al. 2002; Katoh and Standley 2013), and we performed edge trimming on the resulting alignments. Final filtering and concatenation of UCE loci generated matrices of 1031 (60%), 979 (70%), 763 (80%), and 198 (90%) loci. The matrix with ≥ 90% proportion of taxa was selected for use in downstream analyses as support did not decrease with the reduction of included loci. It recovered UCE contigs with a mean length of 532 bp and 122 informative sites.

We used the sliding-window approach and entropy site characteristic (SWSC-EN), a partitioning scheme proposed for UCEs, to determine the best fit scheme for the flanks and cores across each locus in the 90% complete matrix in PartitionFinder 2 (Tagliacollo and Lanfear 2018). We merged the right and left flanks into a single “flank” partition and kept the “core” partition independent. Using ModelFinder implemented in IQ-TREE 1.6.12, we determined the best-fit partitioning scheme (Chernomor et al. 2016; Kalyaanamoorthy et al. 2017; Minh et al. 2020). We used this partitioning scheme to estimate the concatenation-based maximum likelihood (ML) trees in IQ-TREE using 1000 Ultrafast Bootstrap replicates to asses branch support (Hoang et al. 2018). We also conducted coalescent-based species-tree analyses in ASTRAL 5.7.3 using the UCE gene trees as input (Sayyari and Mirarab 2016; Zhang et al. 2018; Rabiee et al. 2019). Gene trees were inferred using the UCE “flank” and “core” partitioning scheme. Given the minimal differences in support between the 70% and 90% complete UCE loci matrices, we used the 90% complete matrix in downstream analyses to reduce computational time (Fig. S1).

### 2.4 Incorporating Public Sequence Data

We included data from eight additional loci, downloaded from either GenBank or assembled from raw sequencing reads, in order to increase taxonomic sampling of the group by including species that did not have genomic data available (Table S1). To extract mitochondrial genes and the nuclear gene *rhod*, we used the pipeline described in Hughes *et al*. (2021). These scripts were additionally modified to include reference sequences from GenBank for the non-coding loci, *12S* (FJ008141.1, EF095566.1, FJ008155.1, LC036890.1) and *16S* (FN688081.1, OM470924.1, EU848456.1). All sequences were aligned using mafft and visually inspected in AliView (Larsson 2014). These resulting alignments were concatenated with the 90% complete UCE alignments into a single supermatrix. We used of partitioning scheme consisting of 22 separate partitions to test differences among the rates of sequence evolution across loci. For protein-coding loci (*ATP6*, *COI*, *COII*, *cytB*, *rhod*, and *nd2*), we assigned each codon position for each gene as a potential partition. For the non-coding loci *12S* and 16S, each locus was considered designated a single partition. To infer the appropriate placement of species where only data from GenBank was available, we used the UCE only ML topology generated previously as a constraint in order to reduce to potential attraction of lineages with missing data. We reran the ML analysis and searched for the best partitioning scheme in IQTREE using the concatenated UCE and exon sequence matrix.

### 2.5 Divergence Time and Rate Estimation

We estimated divergence times of the Mullidae using BEAST 2.6.2 (Drummond et al. 2002; Bouckaert et al. 2014) with multiple calibration points located across the Syngnatharia phylogeny. These included a secondary calibration and six fossil calibrations utilized in Santaquiteria *et al*. (2021), in addition to two fossils that provide higher resolution within our clade of interest (Appendix A). This includes the recently described crown callionymoid fossil species *Gilmourella minuta,* which is estimated to be ∼49 Ma from Monte Bolca (Carnevale and Bannikov 2019). We also included a fossil placed within the modern genus *Mullus*, which is estimated to ∼13 Ma (Carnevale et al. 2006). Given our interest in the timing of biogeographic events in future studies, we did not include any geographical/geminate calibrations within this study. To reduce analysis runtime, we used a restricted subset of the outgroup topology by including only the most divergent pairs of lineages for each calibration node. The only secondary calibration used in this study was located at the root of Syngnatharia, based on ages obtained from other phylogenetic studies (Santaquiteria et al. 2021). Following Santaquiteria et al, we used a uniform distribution with a hard lower bound at 92 Ma and a hard upper bound at 103.5 Ma. All other calibration points used a log normal distribution, and their associated priors are defined in Appendix A.

To estimate divergence times, we used a relaxed clock of log normal distributed rates in BEAST 2.6.2 (Drummond et al. 2002; Bouckaert et al. 2014). A standard birth-death rate model was used as a tree prior. We partitioned the concatenated matrix containing UCE and GenBank data into eleven separate partitions: UCE flank, UCE core, and nine additional partitions based on the best fit partitioning scheme inferred from the initial ModelFinder search (Chernomor et al. 2016). Each partition was given their own unlinked substitution model parameters (Supplemental Material). We used the topology inferred from the ML analyses as a starting topology and we constrained the monophyly of each genus within Mullidae to reduce the erroneous placement of lineages with limited data. Tracer 1.7.1 was used to assess the convergence of two independent analyses of 300 million generations each and confirm proper mixing of all model parameters used in the final Markov Chain Monte Carlo (Rambaut et al. 2018). We used LogCombiner 2.6.3 to both combine these runs and generate a manageable tree set using a 90% burn-in. TreeAnnotator 2.6.3 was used to calculate the maximum clade credibility tree (MCCT) from the tree set, as well as to estimate posterior probabilities at each node. This time calibrated phylogeny was used in subsequent diversification analyses.

### 2.7 Geometric Morphometrics

We performed a morphometric analysis to quantify the variation in body shape across species within the Mullidae. We collected 2D lateral specimen photographs from museum collections, personal databases, primary literature, and aquarium trade, resulting in a total of 253 individuals across 6 genera within Mullidae (Appendix C). A total of 26 landmarks were placed along the specimen image using the R package StereoMorph 1.6.4 (Olsen and Westneat 2015; Fig. 1). To adjust several specimens with preservation artifacts, such as jaws in the open position, we digitally and interactively rotated the jaw tip landmarks around a reconstructed jaw joint position to a closed orientation, using custom trigonometric code, while maintaining a fixed distance, base position and maintaining all other coordinates (Appendix B).

**Fig. 1.**
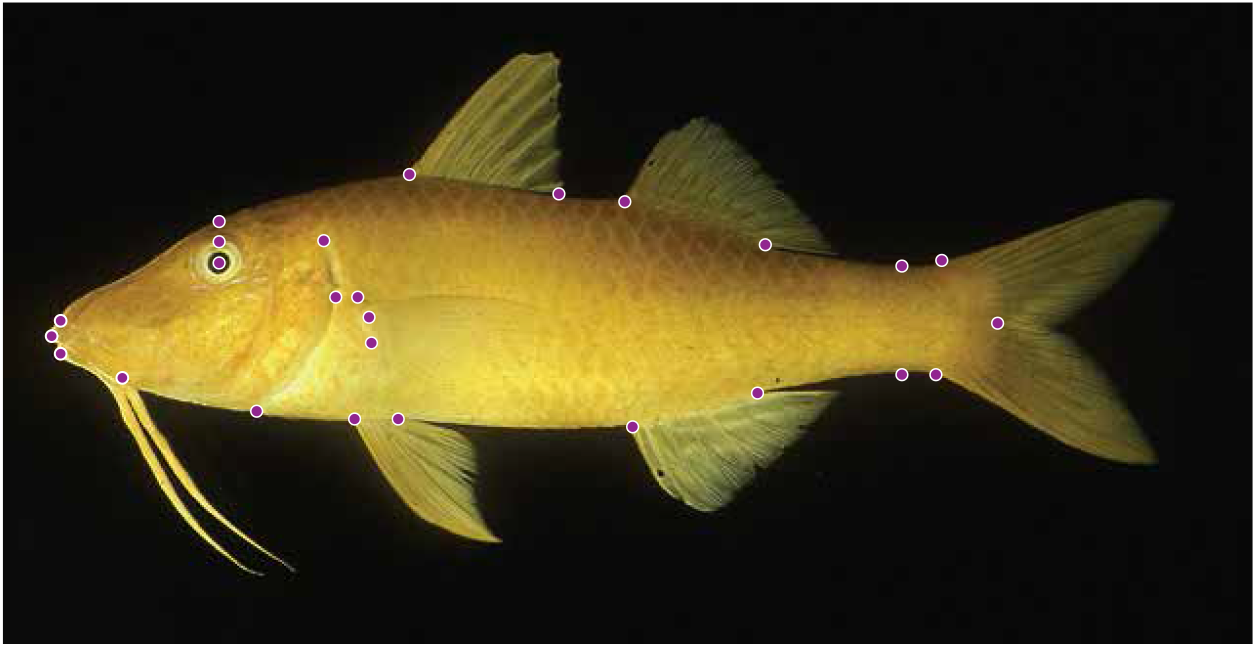
Morphometric protocol for the goatfishes. Full body landmark scheme for the analysis of the geometric morphometrics of goatfishes. The placement location of 26 landmarks with anatomical descriptions are listed in Appendix B.

Using the protocol of George and Westneat (2018) as a general framework, we projected each digitized landmark into linear tangent space using the *gpagen* function in geomorph 4.0.2 (Adams and Otárola-Castillo 2013; Adams et al. 2013, 2021; Baken et al. 2021). The generalized Procrustes analysis (GPA) removes the variation in landmark position that is attributable to rotation, translation, and scaling, while preserving the relevant shape information (Gower 1975; Rohlf and Slice 1990). We then calculated species means of the Procrustes-transformed landmark coordinates to account for intraspecific variation using the *mshape* function in geomorph 4.0.1 (Adams et al. 2021; Baken et al. 2021). This protocol was iterated over all species-grouped landmark sets producing 70 shapes representing the mean body shape of each individual species. From these 70 mean shapes, another GPA was performed and the resulting Procrustes aligned shape data was projected into linear tangent space using the *gpagen* function in geomorph 4.0.1 (Adams et al. 2021; Baken et al. 2021). Principal components analyses (PCA) were performed on the species averaged, Procrustes aligned landmark coordinates using the *gm.prcomp* function in geomorph 4.0.1 (Adams et al. 2021; Baken et al. 2021). We used back transformations to visualize the shape changes associated with the most significant axes of variation (PC1 and PC2), which plots the shape variation in its relative position in the morphospace (Olsen 2017).

In order to assess the evolutionary directionality of shape changes among species, we projected the time-calibrated phylogeny onto the morphospace using the *phylomorphospace* function in the R package phytools 0.7-90 (Revell 2012). Additionally, we generated a phylogenetic PCA using GLS-centering and projection using the implementation in *gm.prcomp* function in geomorph 4.0.1 (Revell 2009; Adams and Otárola-Castillo 2013). To examine the relationship between body shape and genera, we performed a MANOVA using the *procD.lm* function in geomorph 4.0.1 (Adams and Otárola-Castillo 2013). We used the broken-stick method to inform how many principal components (PC) axes to retain for subsequent analyses (Barton and David 1959; Frontier 1976).

The partial disparity for genera was calculated using the overall mean over 1000 iterations with the function *morphol.disparity* in geomorph, taking care to correct the denominator in the variance calculation from n to N-1, where n is the group size and N is the number of observations (Adams et al. 2021; Baken et al. 2021). Procrustes partial variances were reported for each genus and these values were used to calculate the proportion of the total disparity accounted for by each genus (Table S3). We calculated the average disparity through time (DTT) and the mean disparity index (MDI) using the *dtt* function in geiger 2.0.7 (Harmon et al. 2008). Significance was determined using 10,000 null disparity through time simulations generated using the assumptions of Brownian Motion (BM). Additionally, we computed the weight average times, commonly referred to as the Center of Gravity, to examine trends within the distribution of disparity across subclades and time (Slater 2022).

### 2.8 Rates and Pattern of Lineage Diversification

To test hypotheses about the relative rate of diversification across the phylogeny, we first tested deviation from a constant-rate pure-birth diversification process using the γ-statistic (Pybus and Harvey 2000). We controlled for incomplete taxon sampling using the Monte Carlo constant rates (MCCR) test with 10,000 simulations and a sampling fraction of 74.2% (72 out of 102 species), implemented using the *mccr* function in phytools 0.7-90 (Revell 2012). We compared the observed lineage through time (LTT) plot and γ-statistic to the LTT and γ-statistics of 10,000 simulated phylogenies generated under a pure-birth process using the *pbtree* function in phytools 0.7-90 to test for significance (Revell 2012).

## 3. Results

The central product of this research is a well-resolved, time calibrated phylogenetic hypothesis for the Mullidae. The phylogeny is largely congruent with previous taxonomy, and reveals that the crown age of the family is estimated at 21.9 Ma, providing a framework for comparative analysis. Morphometric analysis of body shape strongly supports the current taxonomy and has high phylogenetic signal. Our results reveal that goatfish evolutionary history is characterized by unstable rates of lineage diversification and body shape variation, with a recent increase in lineage accumulation across all genera in just the past 5 million years.

### 3.1 Phylogenomic Analyses and Divergence Time Estimation

The phylogeny of the goatfishes (Syngnatharia: Mullidae) is well resolved and highly supported (Fig. 2 & 3). We find strong support for a monophyletic group with the six described genera: *Mulloidichthys, Mullus, Pseudupeneus, Parupeneus, Upeneichthys and Upeneus*. The topologies derived from the ML, ASTRAL, and time calibrated BEAST analyses were well supported and species relationships, for the most part, were concordant. The Mullidae are sister to the family Callionymidae, which combined represents the suborder Callionymodei within the order Syngnatharia (Fig. 2, Betancur et al. 2017). The divergence between the Mullidae and Callionymidae is estimated to have occurred around 58.2 Ma (48.6 – 74.4 Ma), which resulted in a relatively long branch separating these two families (Fig. 2).

**Fig. 2.**
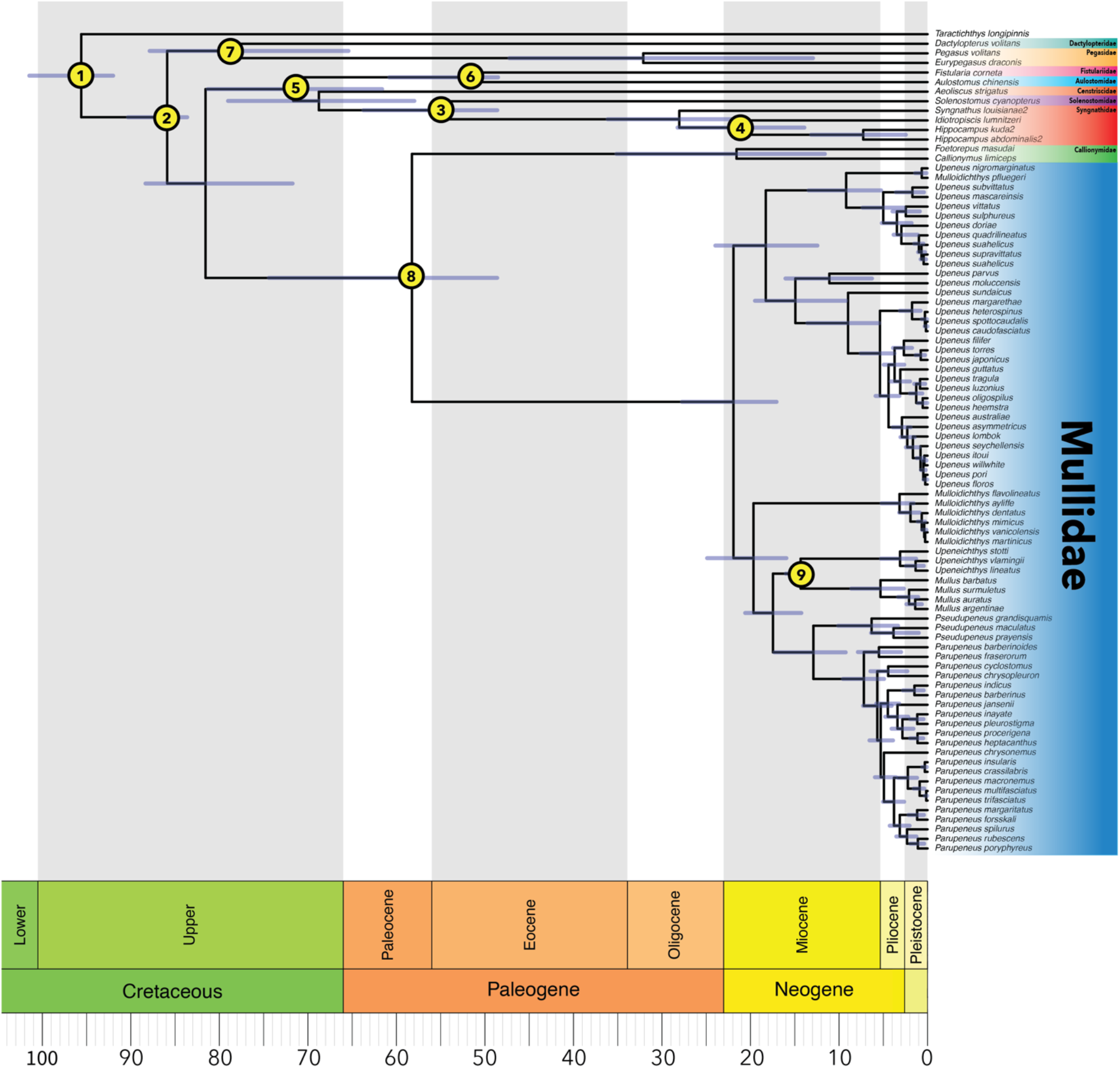
Time-calibrated phylogenetic tree for goatfishes (Family Mullidae) with outgroups. Divergence time estimation for the Mullidae in the context of the outgroups from the order Syngnatharia based on relaxed clock node calibration in BEAST 2. The placement of nine fossil calibrations are shown, and additional information for each calibration can be found in Appendix 1. Blue error bars at each node represent the high posterior density (HPD) of each node age estimate.

**Fig. 3.**
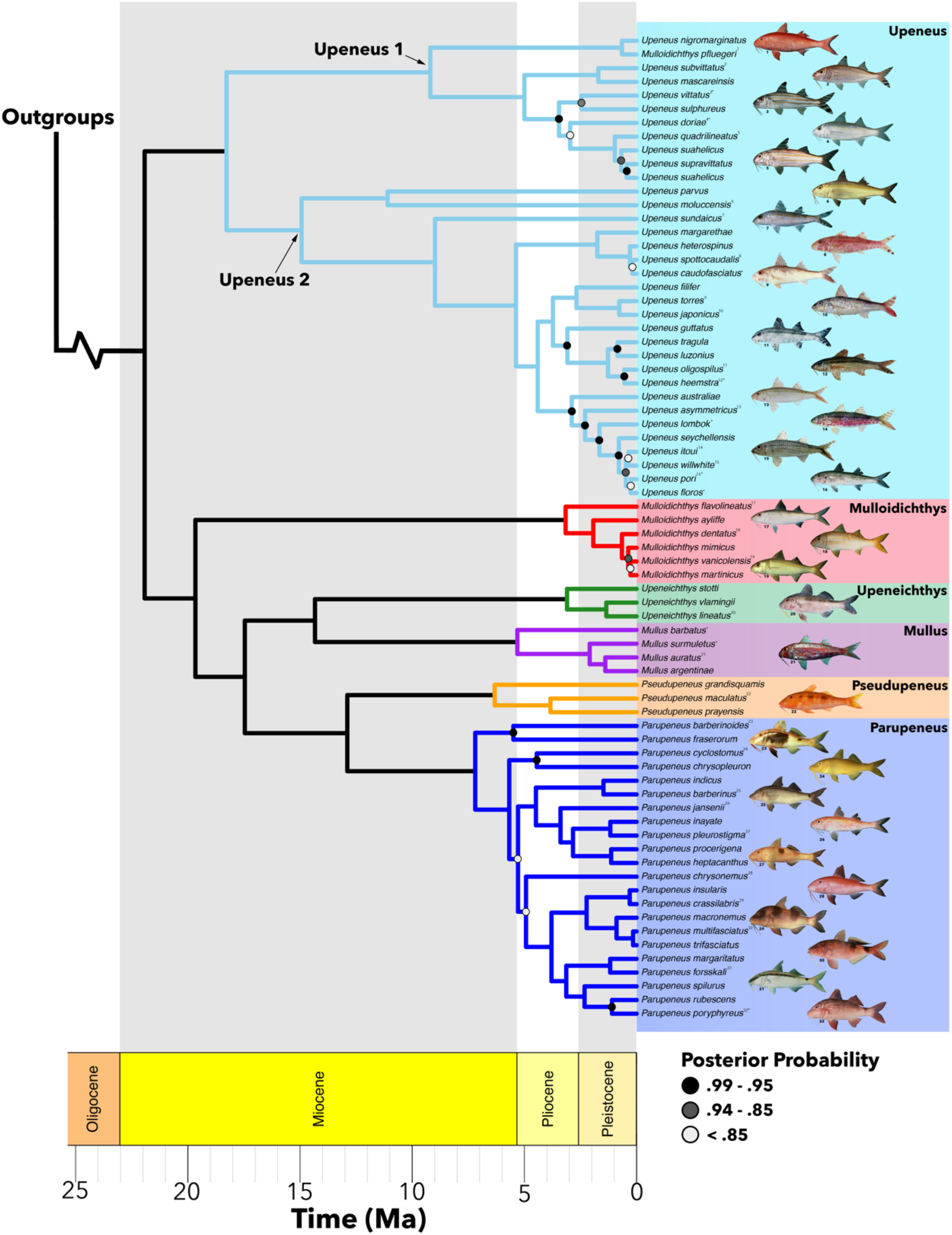
Time calibrated phylogenetic tree for the Mullidae. Divergence time estimates are shown for clades within the Mullidae. Subgenera within *Upeneus* are indicated at each respective node. Nodes without a circle have posterior probability of 1.0. Tips indicated with an asterisk were analyzed with multi-locus data from GenBank only. (Image credits: John E. Randall via FishBase or referenced by number in Appendix 3).

Within the Mullidae, there are two major clades, which split at the root approximately 21.9 Ma (17.0 – 27.7 Ma; Fig. 3). The first clade is comprised of the paraphyletic genus *Upeneus,* which is subdivided into two subclades. The second major clade within the Mullidae is comprised of five genera: *Mulloidichthys, Upeneichthys, Mullus, Pseudupeneus,* and *Parupeneus*. All genera are monophyletic and relationships are well resolved within this clade. There are two main generic sister pairs in this clade, *Mullus + Upeneichthys* and *Pseudupeneus + Parupeneus.* Combined, these genera form monophyletic group with the genus *Mulloidichthys* (Fig. 3).

*Upeneus* is the most species rich genus with 25 species. There are two subclades within the *Upeneus* that diverged approximately 18.3 Ma, denoted as Upeneus 1 and Upeneus 2 (Fig. 3). The only taxonomic anomaly in this clade is the inclusion of *Mulloidichthys pfleugeri,* which is most closely related to the newly described species goatfish species, *Upeneus nigromarginatus* (Bos 2014). A comparison of COI from *M. pfleugeri* across the BOLD database indicates a correct species ID for this sample, and this node has a 100% UF bootstrap support and 1.0 Posterior probability in the ML and ASTRAL analyses, respectively (Fig. 4). Within *Upeneus*, there are four identifiable species group, which we define as a clade of three or more species that diverged within the Pleistocene (Fig. S2). There is one species group in Upeneus 1, which is composed of *U. quadrilineatus, U.suahelicus (taeniopterus), U. supravittatus, and U. suahelicus.* The remaining three species groups are found in Upeneus 2. Group 2.1 is composed of *U. margarethae, U. heterospinus, U. spottocaudalis,* and *U. caudofasciatus.* Group 2.2 is composed of *U. tragula, U. luzonius, U. oligospilus,* and *U. heemstra.* Group 2.3 is composed of *U. asymmetricus, U. lombok, U. seychellensis, U. itoui, U. willwhite, U. pori,* and *U. floros.* (Fig. S2).

**Fig. 4.**
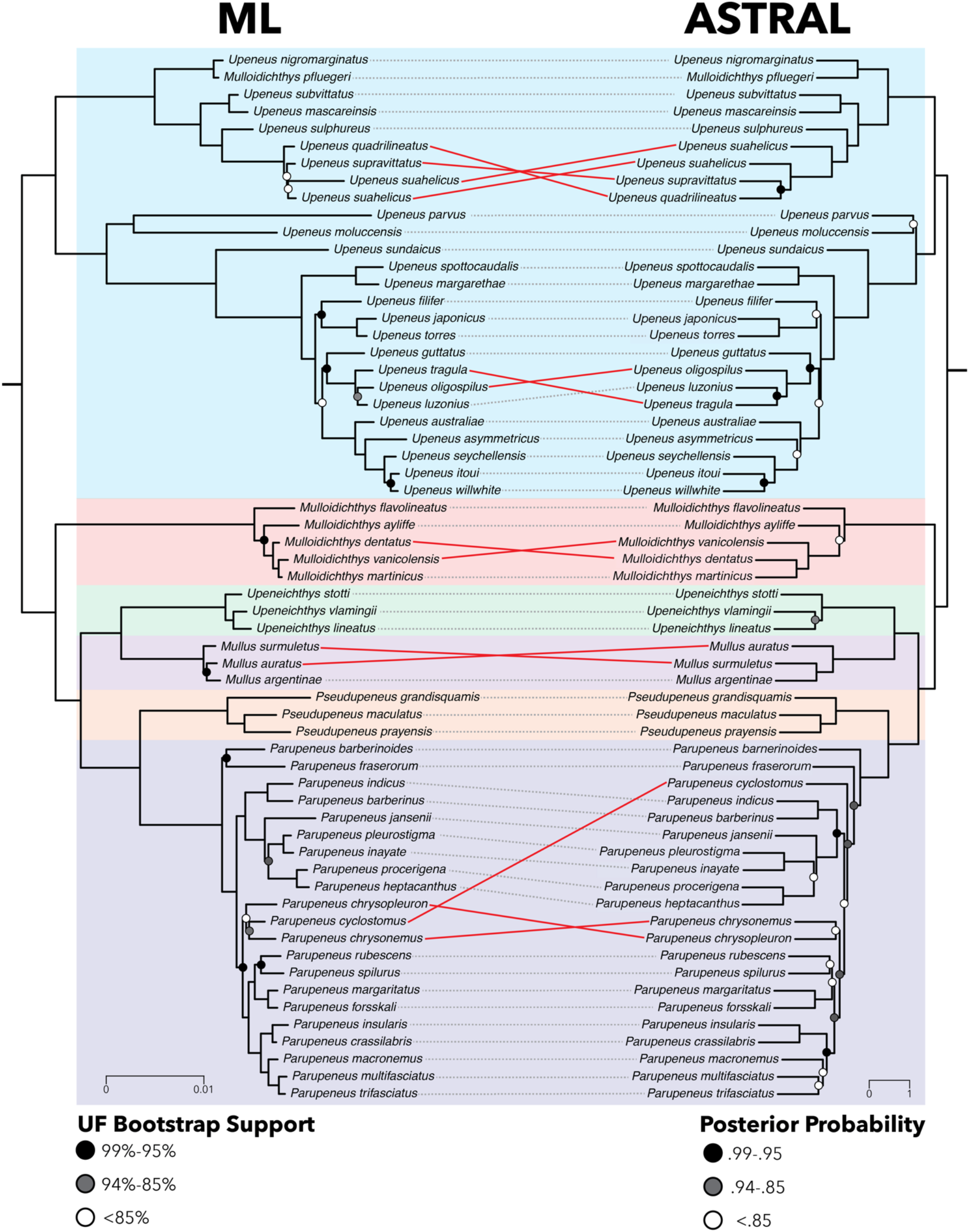
Phylogenetic tree comparisons for the Mullidae. Congruence between the ML tree (left) and the tree inferred using ASTRAL III (right) using the 90% complete UCE matrix. The topologies are largely congruent, with exceptions indicated by a red line between corresponding tips. Nodes without a circle have either 100% Ultrafast Bootstrap support (ML) or a local posterior probability of 1.0 (ASTRAL).

*Parupeneus* is the second most species rich clade with 22 species, followed by *Mulloidichthys* (6 sp.), *Mullus* (4 sp.), *Pseudupeneus* (3 sp.), and *Upeneichthys* (3 sp.). The root age of *Parupeneus* was estimated to be ∼7.1 Ma and the estimated divergence time from its sister genus, *Pseudupeneus,* was ∼12.9 Ma (Fig. 3). There is evidence of recent divergence across *Parupeneus*, with 16 species having diverged within the Pleistocene. However, only two clades met the criteria of species group as the majority of these speciation events were among species pair. The first group is composed of *P. insularis, P. crassilbaris, P. macronemus, P. multifasciatus,* and *P. trifasciatus.* The second group is comprised of *P. spilurus, P. rubescens,* and *P. poryphyreus*.

The mean crown ages of the remaining genera were estimated to be between 6.3 Ma and 3.1 Ma (Fig. 3). When considering *Mulloidichthys, Mullus, Pseudupeneus,* and *Upeneichthys,* the mean difference between root age and stem age is estimated to be ∼9.4 Ma. *Mulloidichthys* has the maximum distance between the root age with a value of ∼9.2 Ma and *Pseudupeneus* has the minimum distance with ∼6.6 Ma between the root age and its divergence from *Parupeneus* (Fig. 3). Similar to the diversification patterns observed in *Upeneus* and *Paruepenus*, the majority of divergence events among species occurred within the middle Pliocene into the Pleistocene. The majority of species within *Mulloidichthys*, with the exception of *M. flavolineatus*, diverged within the Pleistocene and can be classified as a species group. There is a similar pattern of diversification within the majority of species within *Mullus,* consisting of *Mu. surmuletus, Mu. auratus,* and *Mu. argentinae* represent a species group, with the exception of *Mu. barbatus*.

### 3.2 Lineages Through Time

Results of diversification analyses strongly support the hypothesis of unstable rates of diversification throughout goatfish evolution, with a notable inflection point to a sharp increase in lineage accumulation at about 5 Ma (Fig 5A). This pattern of an increase in lineage accumulation is observed within all genera, but is most evident in the most species rich clades, *Upeneus* and *Parupeneus* (Fig. 3). The lineage through time (LTT) plot (Fig. 5A) shows an exponential growth curve with a significant gamma value (γ = 2.4659), which strongly rejects the constant speciation hypothesis under a Yule pure-birth model (p = 0.004). This result indicates that there is a variable rate of diversification across the Mullidae phylogeny, which is evident by a reduction in lineage accumulation during the Middle to Late Miocene prior to the striking acceleration of lineage accumulation during the Miocene-Pliocene transition (Fig. 5A). Additionally, over 50% of the mean node ages are estimated as less than 5 Ma (Fig. 5B), with 92% of total species (66 out of 72 species) divergent from their most recent ancestor within the past 5 Ma.

**Fig. 5.**
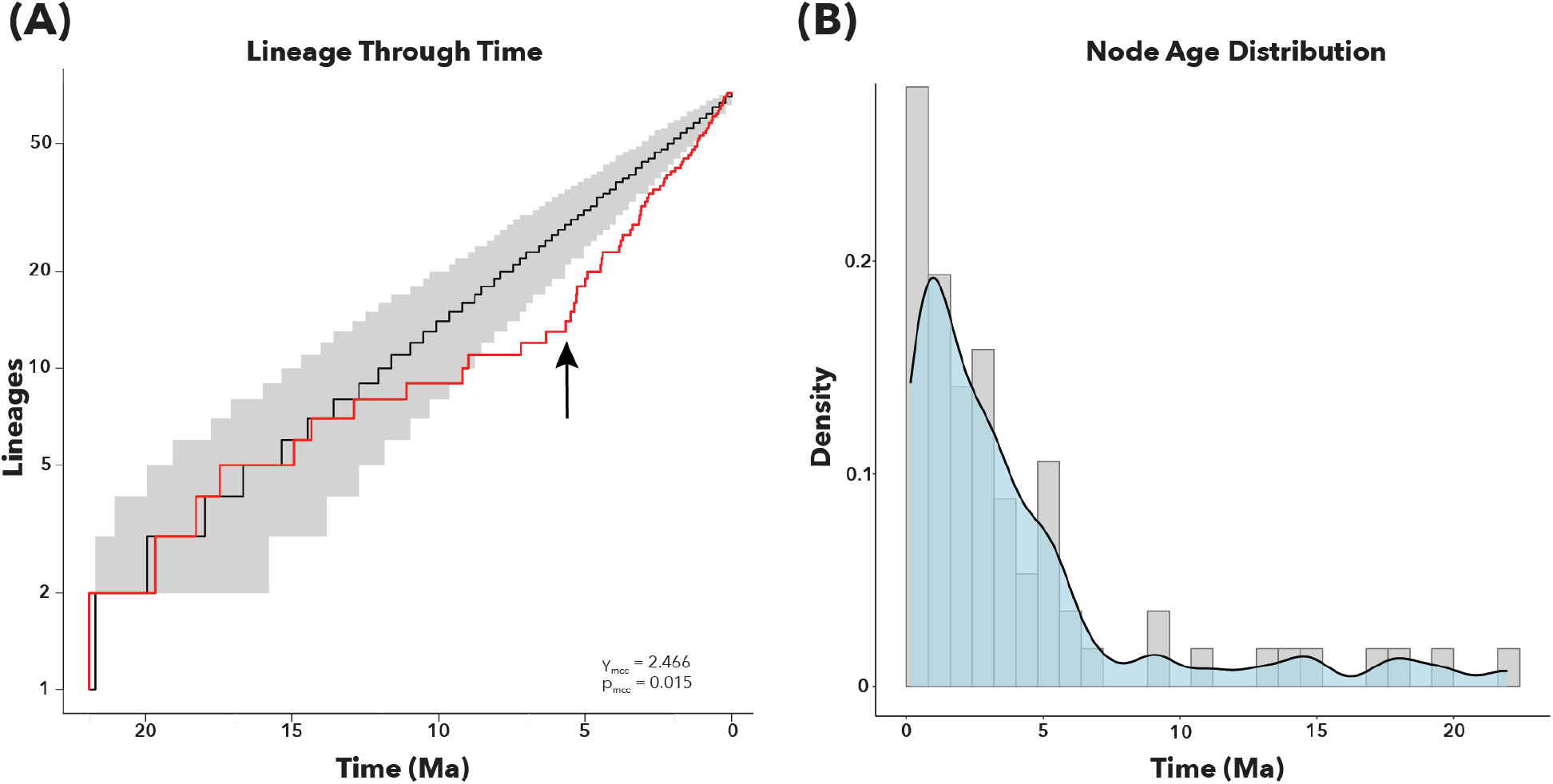
Goatfish diversification patterns through time. **(A)** Trends in lineage accumulation through time. The red curve indicates the data from this study, the black curved indicates the mean of the simulated phylogenies, and the grey area represents the 95% confidence interval of the simulations. Note the inflection point which indicates the increase in lineage accumulation at approximately 5 Ma. (**B)** Density of nodes with ages at specific time intervals.

### 3.3 Body Shape Morphometrics

Body shapes cluster based on genera and have high phylogenetic signal. The primary axis of variation (PC1: 45.26%) for this dataset describes the relative ratio of length to width, termed the elongation ratio (Claverie and Wainwright 2014; Friedman et al. 2019; Fig. 6). Higher PC1 scores are associated with a lower elongation ratio, which is associated with variability on height in the across the body and to a relatively shorter distance between the tip of the mouth and edge of the caudle peduncle. This is especially evident when comparing the height and location of the base of the first dorsal fin to the height of the caudle peduncle, for example between *Parupeneus* and *Upeneus* (Fig. 6). Lower PC1 scores are associated with a higher elongation ratio, which corresponds to species with longer, more evenly slender body shapes. The second axis of variation (Fig. 6; PC2: 14.11%) is associated with eye size, the relative distance between the bases of the second dorsal fin and the caudle peduncle, and body depth in relation to the midline (Fig. 7). Principal components analysis shows that 90% of the variance in body shape is summarized within PC1 through PC10.

**Fig. 6.**
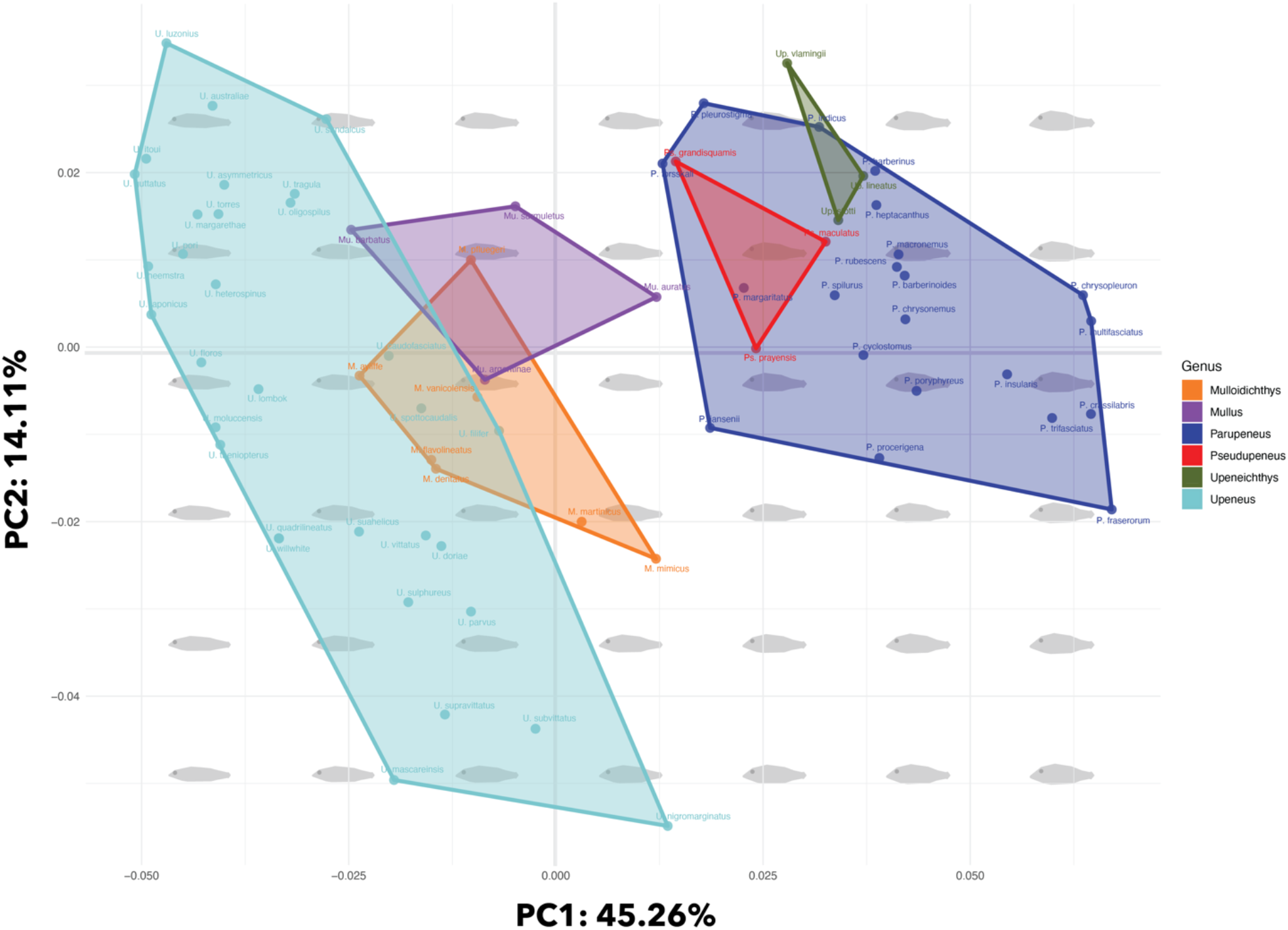
Morphospace for the body shape of the goatfishes (Family Mullidae). Axis 1 and 2 from Principal Component Analysis (PCA) on Procrustes aligned body shape landmarks for 70 goatfish species. Each coordinate represents the mean shape of each species. PC1 represents 45.26% of the total variation, and PC2 represents 14.11% of the total variation. Species are grouped by genus, and this is indicated by color coded hulls to show the distribution of each genus in morphospace. Gray polygons show the back transformation, which represents the hypothetical goatfish body shape at intervals along each axis.

**Fig. 7.**
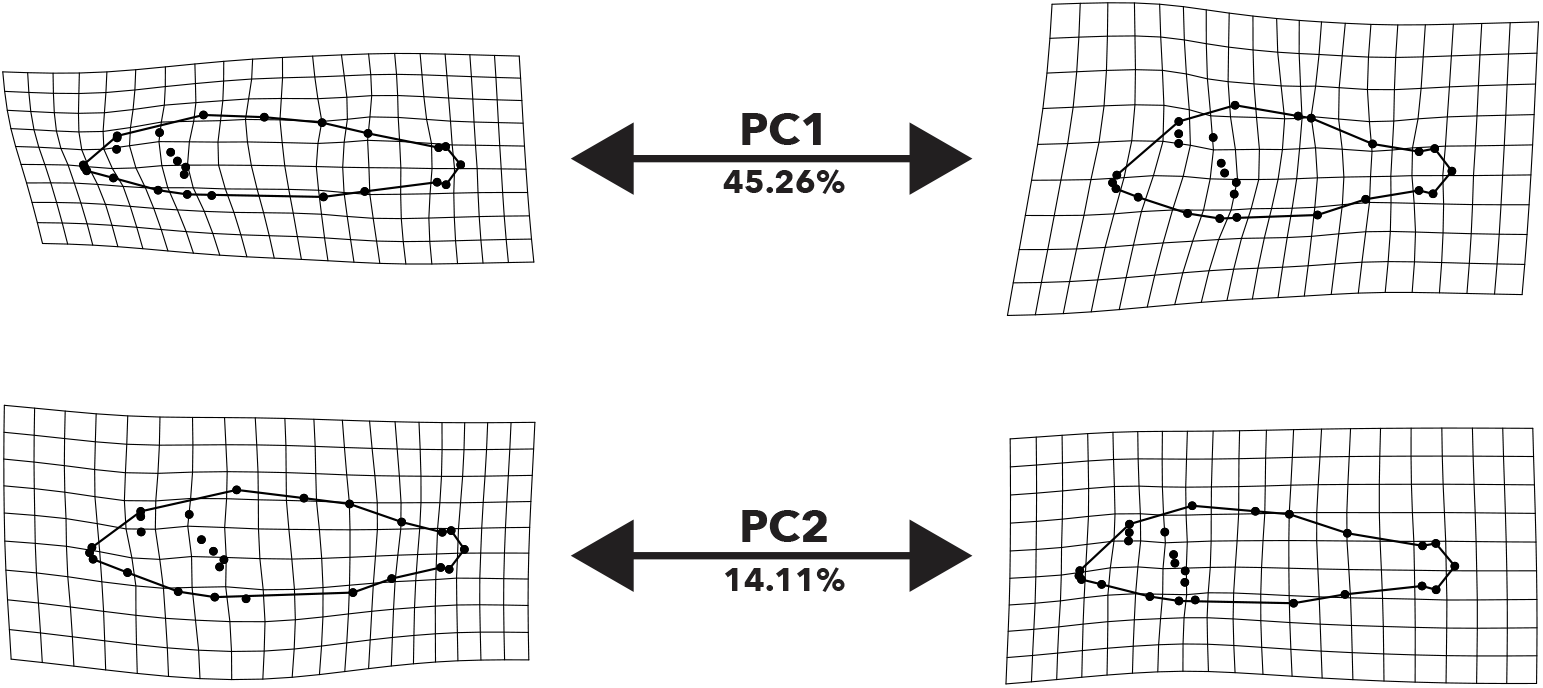
Thin-plate spine warp grids for PC1 and PC2 of goatfish body shape. These plots show the relative deformation associated with the extreme ends of PC1 and PC2 compared to the overall mean shape.

Genera are significantly associated with body shape and segregate primarily along PC1 into two main groups. Group 1 is composed of *Upeneus, Mulloidichthys,* and *Mullus*, and Group 2 consists of *Parupeneus, Pseudupeneus*, and *Upeneichthys*. Group 1 is clustered on lower on the axis of PC1 and Group 2 is clustered on higher values of PC1. Of note, *Pseudupeneus* is fully encapsulated within the *Parupeneus* morphospace. Although genera do not cluster across PC2, there are differences among the ranges of values in which each genus occupies. Species within *Upeneus* span the entirety of PC2, which ranges highest to lowest represented by *U. luzonius* to *U. mascareinsis,* respectively (Fig. 6). PC2 does appear to separate the two subclades within *Upeneus*, with Upeneus 1 occupying higher values of PC2 and Upeneus 2 occupying lower values of PC2 (Fig. S4). However, these groups are not fully distinct as there is overlap within values of PC1 (Fig. S4). The only other genus that has species that falls within similarly high ranges of PC2 is *Upeneichthys*, represented by *Up. vlamingii*. There are no species that have both high values of PC1 and low values of PC2, which is represented by the empty lower right quadrant in the morphospace. *Upeneus* and *Parupeneus* have the highest proportion of total variance (Table S3) and a MDI of 0.098.

### 3.4 Body Shape Morphometrics Across Phylogeny

Analyses revealed a significant phylogenetic signal in the body shapes of goatfishes. The phylomorphospace estimates the root node within the lower left quadrant, indicating low values of PC1 and PC2. When considering phylogenetic relationships among the genera, the two morphotype groups are largely congruent with phylogenetic structure (Fig. 8A & S4). The main exception to this pattern is the convergence of *Upeneichthys* with *Parupeneus* in morphospace, despite its sister genus being *Mullus* (Fig. 8A). When accounting for the variance that is correlated with phylogenetic relationships, there is a strong overlap among the morphotype groups (Fig. S5). All genera were centered around the origin, although there were differences between the distribution of the two most species rich genera, *Parupeneus* and *Upeneus*. *Parupeneus* and *Upeneus* have equivalent variance across PC1, and *Parupeneus* and *Upeneus* 1 split with *Upeneus* 2 along PC2. There is a significant deviation from the null expectations for disparity through time (Fig. 8B & S6). Additionally, there is relatively late maintenance of higher within subclade variation than we would expect with BM into the Pliocene and Pleistocene (Fig. 8B & S6).

**Fig. 8.**
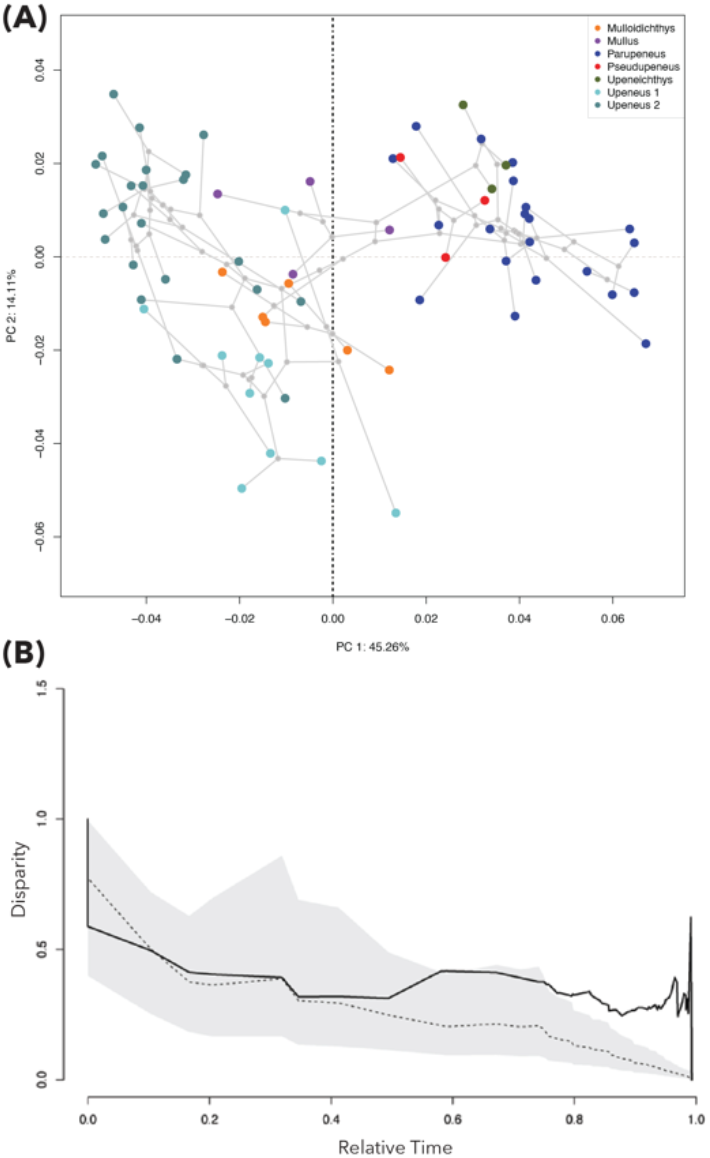
Phylomorphospace and body shape variability through time for the goatfishes. **(A)** Phylomorphospace projections of body shape. Each tip is colored according to clade membership. Grey branches indicate the internal branch structure of the superimposed phylogeny shown in Figure 3. (**B)** Disparity Through Time (DTT) plots for PC1 – PC3 of body shape. Disparity is the average subclade disparity divided by the total clade disparity for each internal node of the tree. The solid black curve indicates the trait disparity values for this study, and the dashed curve is trait disparity for simulated data under Brownian Motion. Grey shaded areas of each plot indicate the 95% confidence interval inferred from the simulations.

## 4. Discussion

The addition of a robust phylogeny of goatfishes to the growing marine fish Tree of Life enriches our understanding of the tempo and mode of coral reef fish biodiversity and evolution. The central conclusions of this study are 1) phylogenetic relationships among the species and genera of Mullidae were highly resolved, 2) body shape morphometrics revealed several diverse body profiles among the goatfishes, largely congruent with phylogeny and taxonomy position, and 3) we found strong evidence for unstable rates of phylogenetic and morphological diversification, with an unusual pattern of recent increase in lineage accumulation across all genera in the family that may be associated with habitat changes at the Miocene-Pliocene transition.

### 4.1 Highly supported phylogenetic relationships within goatfishes

Our phylogenetic analysis represents the most species rich Mullidae phylogeny to date with the inclusion of 72 species, representing just over 70% of extant goatfishes. We resolved the Mullidae into six highly supported genera, which were monophyletic with the exception of *Mulloidichthys* (Fig. 3). The majority of internal nodes were well supported and the phylogenetic relationships were largely consistent across inferences made using ML, ASTRAL, and Bayesian approaches with only a few notable exceptions (Fig. 3 & 4). We found evidence that some species groups have very short branch lengths between taxa, which made it difficult to accurately resolve the tip level relationships among some species. This is most evident with the species group in the *Upeneus* 1 clade, comprised of *U. quadrilineatus, U. suahelicus (taeniopterus), U. supravittatus, and U. suahelicus.* The branch lengths between these species are extremely short, resulting in different topologies with low support across all three of our inference methods (Fig. 3 & 4). Another interesting incongruence is *Parupeneus cyclostomus*, which has a large shift in its phylogenetic placement that is particularly evident when comparing the ML and ASTRAL topologies (Fig. 4). This is again likely the result of short branch lengths that result in low phylogenetic support and low gene tree concordance among *P. cylcostomus, P. chrysonemus,* and *P. chrysopleuron* (Fig. 4).

While our phylogenetic analyses generally support the results of previous studies, there are several main exceptions that can be found across the goatfish topology. In comparison with the clade-wide phylogeny inferred from solely morphological characters, our results resolve a slightly different relationship structure among genera with strong support for the sister relationship of *Mullus* and *Upeneichthys* (Kim 2002). Additionally, our inferences resolved many of the polytomies found in that study, primarily among closely related species. Interestingly, many of the phylogenetically informative morphological synapomorphies of each genus were well supported when examining the goatfishes using genomic data. The other primary source of phylogenetic incongruence with previous studies was found when examining the relationships among species within closely related species complexes, primarily within *Upeneus* and *Parupeneus* (Fig. S7). These species complexes are often described based on similarities taxonomic traits, such as the number of fin rays and geographic proximity (Uiblein et al. 1998; Uiblein and Heemstra 2010, 2011; Uiblein and Gouws 2015; Uiblein and Motomura 2021). It is important to note that there is uncertainty across many of the nodes associated with species that are assigned to species groups, which reaffirms the issue of resolving the relationships among species separated by very short branch lengths (Fig. 4 & S7). Despite this, the majority of described species complexes cluster in morphospace, which indicates some degree of similarity across body shapes (Fig. S8).

### 4.2 Recent Diversification of Goatfishes

Recent genomic studies on the order Syngnatharia have estimated a far younger crown age of the goatfishes than previously suggested (Betancur et al. 2017; Santaquiteria et al. 2021). The first study to include an estimation of the crown age of the goatfishes was Near et al. (2013), which estimated an age of approximately 40 Ma based on ten nuclear genes for five species (Near et al. 2013). Subsequent studies resulted in a large amount of variation in the estimated crown age, with ages ranging from 20 Ma to 67 Ma depending on the type of genomic data used and the number of species included (Betancur-R et al. 2013; Betancur et al. 2017; Alfaro et al. 2018; Hughes et al. 2018; Rabosky et al. 2018).

In this study, we estimated the crown age of goatfishes to be approximately 21.9 Ma (HPD 17.0 – 27.7 Ma) and the divergence between Mullidae and Callionymidae to have occurred approximately 58.2 Ma (HPD 48.6 – 74.4 Ma; Fig. 2). Our results most closely resemble the findings of Santaquiteria *et al*. (2021), which estimated a slightly younger mean age of goatfishes at 18.0 Ma and high probability density (HPD) node age range of 14.9–21.8 Ma (Santaquiteria et al. 2021). However, our estimated divergence time between Mullidae and Callionymoidei is much younger than Santaquiteria et al. (2021), as their estimates date the split during the Late Cretaceous (Santaquiteria et al. 2021). This variation in the observed HPD ranges and outgroup divergence times could be due to diffferences in the calibration scheme. In this study, we did not use any germinate calibrations to constrain the node ages based on geologic ages, and we included two additional fossil calibrations located at the stem node and and an internal node within the Mullidae (Fig. 2). For example, we included a fossil *Mullus* from Northern Russsia dated as approximatly 13 Ma, which is relatively close in age to the lower age estimate for the root of Mullidae in Santaquiteria *et al*. (2021). Additionally, our inclusion of more species in the crown group could push the age range older due to the higher resolution of species level relationships.

The estimated crown age of goatfishes is notably younger than many comparable marine fish clades of this size and geographic distribution (Tedesco et al. 2017; Alfaro et al. 2018; Miller et al. 2018; Rabosky et al. 2018). For example, the majority of modern coral reef associated fish families have fossil representatives in the early Eocene deposits at Monte Bolca, with only butterflyfishes originating after the Eocene at approximately 32 Ma (Cowman and Bellwood 2011; Bannikov 2014; Bellwood et al. 2017). This young origin age is evident even among comparisons with the other suborder level clades within the Syngnatharia, as all of the other major suborder clades within the Syngnatharia diverged within the Late Cretaceous (∼70-83 Ma; Santaquiteria et al. 2021). These divergence ages are drastically older than the estimated age of the Mulloidei, a suborder that is comprised of the Mullidae (Fig. 2).

Additionally, our inferences resulted in a relatively long branch length between the stem and crown node of the goatfishes, which spans from the late Oligocene to the early Miocene (Fig. 2). The Oligocene – Miocene transition was characterized by a relative decrease in global temperature and fluctuations in Antarctic ice volume, as well as significant changes in marine biogeographic regions, which resulted in the migration of the warm water faunas of the Indo - Pacific into the Mediterranean basin (Beddow et al. 2016). Given the limited availability of fossil data within the goatfishes, it is impossible to accurately estimate possible extinction events, and therefore diversification, along the stem branch (Louca and Pennell 2020). However, this transition marked the first major divergence at the root of the crown goatfishes as the circumglobally distributed genus *Upeneus* split from the remaining genera (Fig. 2 & 3).

### 4.3 Morphological differentiation among genera

The geometric morphometrics of body shape among species within the Mullidae highly supports the phylogenetic placement and previous taxonomy. Our analyses reveal two main morphotypes that diverged along PC1, along which variation is primarily associated with differences in body elongation, head depth, and fin base position (Fig. 6 & 7). Each of these morphotypes are significantly associated with a particular set of genera. For example, morphotype 1 is comprised of *Upeneus, Mulloidichthys,* and *Mullus* which tend to be slender and more elongated. In contrast, morphotype 2 is comprised of *Upeneichthys, Pseudupeneus,* and *Parupeneus* and is associated with truncated body length and a larger head depth. These results are consistent with findings from previous studies on body shape evolution across fishes, which demonstrated that the major axis of variation was often associated with body elongation (Claverie and Wainwright 2014). Body elongation, often described as elongation ratio, has many functional implications and may be associated with habitat, trophic mode, and swimming performance across a wide variety of fish clades (Claverie and Wainwright 2014; Friedman et al. 2019). Another important aspect of morphological variation associated with PC1 is head shape. Head shape is often associated with trophic specialization, as the feeding mechanics that underlie head morphology often impose restrictions on the types and range of possible morphological variation (i.e. Cooper and Westneat 2009; López-Fernández et al. 2013; Mahe et al. 2014). Variation along PC2 is associated with eye diameter, ventral body depth, and the relative distance between the base of the second dorsal fin and the caudle peduncle (Fig. 6 & 7). Variation across these traits is associated primarily with habitat and prey capture. Interestingly, only species within *Upeneus* occupy the portion of morphospace associated with lower values of PC2 (Fig. 6).

All species of goatfishes are specialized benthic carnivores that use their tactile and chemosensitive barbels to disturb the substratum in search of food (Gosline 1985; McCormick 1993, 1995; Platell et al. 1998; Lukoschek and McCormick 2001). Despite their relative similarities in feeding morphology, previous research has shown that co-occurring species often differ in their substrate preferences, feeding modes, and diet compositions (Gosline 1985; Golani 1994; McCormick 1995; Platell et al. 1998; Krajewski et al. 2006). Given that the precise relationships among head and body shape, substrate preference, and feeding mode are currently unknown across all species of goatfishes, future work should test the functional and ecomorphological associations among these traits.

Our results show a highly significant association among genera, body shape, and head shape. Divergence in body and head shape is consistent with a BM model of evolution, in which lineages exploited new regions of morphospace during the early divergences among genera (Fig. 8a). However, the distribution of PC scores across the phylogeny suggests that morphotype 2 is derived from morphotype 1, and may be associated with changes in preferred benthic substrates and feeding behaviors (Fig. S3). Interestingly, no lineages occupy the area of morphospace defined by high values of PC1 and low values of PC2, which is associated with truncated body, larger body depth, and larger eye size (Fig. 6). Additionally, we do not find evidence for significant differences in the amount of disparity among genera. These evolutionary patterns suggest that there were early divergent strategies across morphotypes early in goatfish history, and species have subsequently radiated within their distinct areas of morphospace. Further research needs to be done to test hypotheses about potential ecological or functional constraints and whether this is an example of phylogenetic niche conservatism within genera.

### 4.4 Phylogenetic and Morphological Diversification Associated with Miocene-Pliocene Transition

The transition from the late Miocene into the early Pliocene marked noticeable changes in the rates of lineage accumulation across all genera of goatfishes. There was a pulse of rapid diversification beginning approximately 5 Ma, which is evident by the presence of short branch lengths and the high density of nodes with ages less than 5 Ma (Fig. 3 & 4). This pulse follows a period of relatively slower rates of diversification during the middle to late Miocene, which is evident by long branch lengths and significantly lower rates of lineage accumulation (Fig. 3 & 4). Although goatfishes have been examined in the context of large-scale marine diversification studies now (Siqueira et al. 2016; Alfaro et al. 2018; Rabosky et al. 2018), these variable diversification rates have not been observed until now. The timing of this increase in lineage accumulation during the end of the Miocene and into the Pliocene corresponds with the maintenance of greater mean subclade disparity than expected under pure BM, which could indicate that species were simultaneously undergoing changes in the rates of both phylogenetic and morphological diversification (Fig. 4 & 8B). The timing of this variability in diversification rates suggests that the paleogeographic, climatic, and sea level changes associated with the Miocene-Pliocene transition played an important role in the evolution of goatfishes. In particular, the end of the Miocene is associated with an increase in the interchange of Indo-West Pacific fauna caused by the separation of the Arabian and African plates, despite the complete separation of the Mediterranean Sea from the Red Sea and the Indian Ocean due to terrain uplift (Briggs 1995).

The timing of lineage accumulation and morphological diversification most closely resembles the trends observed in studies of adaptive radiations, which often occur in limited geographic areas and in the absence of competition (Glor 2010; Hench et al. 2022; Thacker et al. 2022). Although there is evidence for similar rates of diversification in fish clades such as the parrotfishes, hamlets, and gudgeons, it is unusual for a clade of the species richness and distribution of goatfishes to have recent bursts of speciation across all genera (Smith et al. 2008; Hench et al. 2022; Thacker et al. 2022). One example can be found in a recent study by Knudsen et al. (2019), which observed rapid diversification in the Kyphosidae (sea chubs), during the middle Miocene into the Pleistocene that was likely associated with their generalist herbivorous diet and pelagic larval dispersal (Knudsen et al. 2019). However, the species richness of Kyphosidae is much lower than the goatfishes with only 16 described species despite their circumglobal distribution (Knudsen and Clements 2016; Fricke et al. 2022).

Given our current understanding of goatfish ecology and life history, high rates of larval dispersal and specialized feeding modes likely play a large role in driving recent bursts in diversification. Water column position has been shown to correlate with rates of morphological diversification, as benthic associated species may be associated with higher diversification rates compared to their pelagic counterparts (Friedman et al. 2020; Rincon-Sandoval et al. 2020). However, this trend has been shown to vary widely across clades depending on their developmental patterns and life history traits (Claverie and Wainwright 2014; McCord et al. 2021). Population level studies have shown that species of goatfish have been able to maintain panmictic populations across large geographic scales due to their long pelagic larval duration (Lessios and Robertson 2013; Fernandez-Silva et al. 2015). Interestingly, evidence for population isolation and differentiation was observed primarily at the periphery of the species’ range (Lessios and Robertson 2013; Fernandez-Silva et al. 2015), which suggests that recent geographic changes in habitat availability can impact the ability of species to effectively disperse and subsequently diversify.

### 4.5 Conclusion

A species rich time-calibrated phylogeny of the Mullidae allows us to make important additions to our understanding of goatfish evolution and diversification. Phylogenetic relationships among species and genera are highly resolved, and corroborate with the structure among body shape morphologies. The Miocene-Pliocene boundary marked a period of both rapid changes in the rates of phylogenetic and morphological diversification, which was possibly driven by changes in the availability of suitable habitats caused by geologic changes. An increased understanding of the evolutionary relationships among key groups of organisms will continue to reveal surprising trends in the tempo and mode of diversification that resulted in the patterns of biodiversity that we observe today.

## Supporting information

Table S1 Tissue and Sequence Info

Table S2 Genbank Accession Table

Table S3 Clade Disparity

Appendix A Fossil Calibration Information

Appendix B Morphological Landmarks

Appendix C Photo Sources

## Acknowledgments

We thank the following people and institutions for providing preserved tissues for genomic sequencing: Mark McGrouther, Amanda Hay, and Joey DiBattista from the Australian Museum (AMS), Richard Pyle and Ken Hayes from the Bishop Museum of Natural History (BPBM), Luiz Rocha from the California Academy of Sciences (CAS), Alistair Graham and Will White from CSIRO, Hiroyuki Motomura from the Kagoshima University Museum (KAUM), Michael Berumen from the King Abdullah University of Science and Technology (KAUST), Andy Bentley and Leo Smith from the University of Kansas (KU), Prosanta Chakrabarty from Louisiana State University Museum of Zoology (LSUMZ), and Amanda Gura, Wouter Holleman, and Gavin Gouws from the South African Institute of Aquatic Biodiversity (SAIAB). Additional tissues were generously donated from personal collections by Arthur Bos and Giacomo Bernardi. We especially thank Caleb McMahan, Susan Mochel, and Kevin Swagel from the Field Museum of Natural History (FMNH) for their assistance in tissue storage, preparation, and shipping. Additionally, we thank Rose Peterson (George Washington University) for assisting with DNA extractions. We thank Andrew George (University of Chicago), Aintzane Santaquiteria (University of Oklahoma), Michael Coates (University of Chicago), and Graham Slater (University of Chicago) for their generous and insightful comments throughout the progress of this study. We also thank Cameron Hill (Wesleyan University) for incredibly helpful discussions and advice on morphometrics related statistics and Franz Uiblein (Institute of Marine Research, Norway) for taxonomic input during the review process. We used the Midway2 high performance computing cluster operated by the Research Computing Center at the University of Chicago for all genomic analyses used in this research.

## Data Availability Statement

Genomic data will be made accessible at BioProject PRJNA824087. All other data and files are stored on FigShare, which are accessible through the following links (available when published).

