## Supplemental Information for "Phylogenomics and body shape morphometrics reveal recent diversification in the goatfishes (Syngnatharia: Mullidae)"

**Supplemental Information for “Phylogenomics and body shape morphometrics reveal recent origin and diversification in the goatfishes (Syngnatharia: Mullidae)”**

**Supplemental Figures:**

1. ML phylogeny of the goatfishes inferred using a 70% complete UCE matrix
2. Phylogenetic placement of goatfish species groups
3. Body shape morphospace for goatfishes labeled by clade membership
4. ContMap showing the phylogenetic distribution of PC1 and PC2 scores
5. Phylogenetic PCA of goatfish body shape
6. Center of Gravity distribution
7. Described species groups within the Mullidae
8. Body shape morphospace for the Mullidae labeled by species complex membership

**Supplemental Tables:**

1. Tissue and Sequence Information
2. GenBank Accession Table
3. Proportion of Disparity by Clade

**Appendices:**

1. Fossil Calibration Information
2. Body Shape Landmark Placement and Information
3. Photo Sources

**Files:**

1. **ASTRAL**
   a. astral_script.sh – *bash script for running ASTRAL on the HPC*
   b. mullidae_only_uce_genetrees.trees – *gene trees for each UCE locus*
   c. Mullidae_Only_Astral_Phy.tre – *ASTRAL phylogeny of UCEs*
2. **IQTREE**
   a. Mullidae Only
      i. UCE Only
         1. IQTree_Mullidae_Only_uce_partition_script.sh – *bash script*
         2. Mullidae_Only_uce90_alignment.fasta – *UCE Alignment (90%)*
         3. Mullidae_Only_uce90_partition_best_scheme.nex – *Best partition scheme*
         4. Mullidae_Only_uce_only_ML_PHY.nexus – *ML Phylogeny UCE only*
      ii. Mulidae Combined (UCE + GenBank)
          1. IQTree_MO_combined_script.sh – *bash script*
          2. Mullidae_Only_combined_alignment.fasta – *UCE & GB alignment*
          3. Mullidae_Only_uce_combined_best_partition_scheme.nex – *best partition scheme*
          4. Mullidae_Only_uce_combined.treefile – *ML combined phylogeny*
   b. With Outgroups
      i. UCE Only
         1. IQtree_Mullidae_uce_partition_script.sh – *bash script*
         2. 70% Completeness Matrix
            a. Mullidae_uce70_outgroup_alignment.fasta – *UCE 70% alignment*
            b. Mullidae_uce70_outgroup_MLphy.nex – *ML phylogeny 70%*
         3. 90% Completeness Matrix
            a. Mullidae_uce90_outgroup_alignment.fasta – *UCE alignment 90%*
            b. Mullidae_uce90_outgroup_partition_best_scheme.nex – *best partition scheme*
            c. Mullidae_uce90_outgroup_ML.treefile – *ML phylogeny 90%*
      ii. Combined (UCE + GenBank)
          1. IQTree_Combined_Scripts.sh – *bash script*
          2. Mullidae_All_Genbank_Alignment.fasta – *Genbank alignment with outgroups*
          3. Mullidae_uce90_combined_alignment.fasta – *Combined alignment*
          4. Mullidae_uce90_combined_outgroup_partition_best_scheme.nex – *best partitioning scheme*
          5. Mullidae_uce90_combined_outgroup.treefile – *ML combined phylogeny with outgroups*
3. **BEAST**
   a. mullidae_beast_script.sh – *beast bash script*
   b. Mullidae_beast_input.xml – *beast XML*
   c. Mullidae_beast_input_run1.log – *beast logs from Run 1*
   d. Mullidae_beast_input_run2.log – *beast logs from Run 2*
   e. Mullidae_beast_combined.log – *combined beast log*
   f. Mullidae_beast_combined_b90.trees – *posterior treeset from 90% burn-in*
   g. Mullidae_beast_combined_b90_mcc.trees – *MCC phylogeny from 90% burn-in*
