## Supplemental Figures for "Phylogenomics and body shape morphometrics reveal recent diversification in the goatfishes (Syngnatharia: Mullidae)"

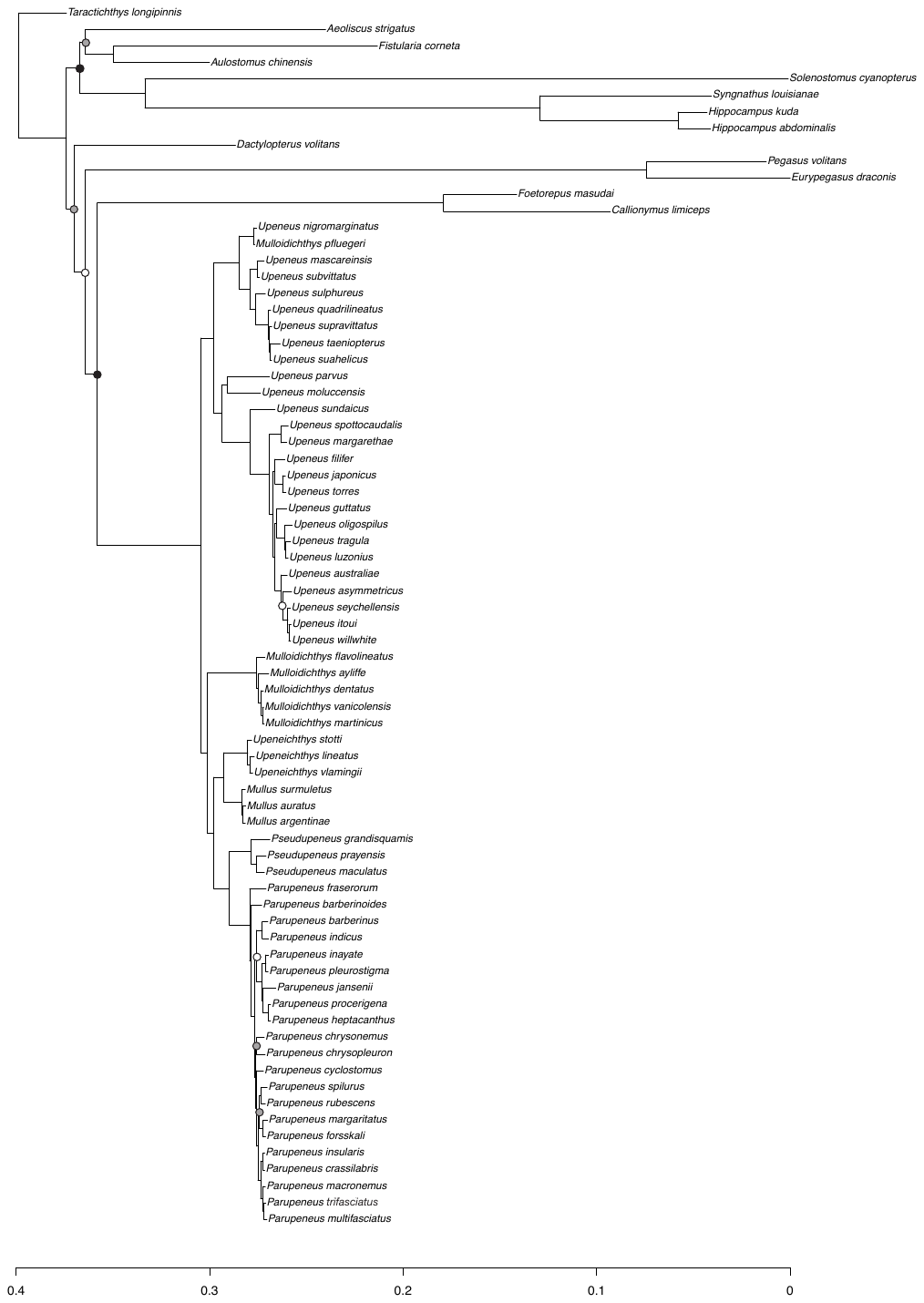

**Supplementary Fig. 1.** Maximum likelihood phylogeny of the goatfishes inferred using a 70% complete UCE matrix in IQTREE. UF bootstrap support values are shown using color coded circles on certain nodes. Nodes without a circle have a UF Bootstrap support of 100%, while nodes shaded black = 99% - 95%, grey = 94%-85%, and white = <85%. The relevant tree and alignment files can be found in supplementary data.

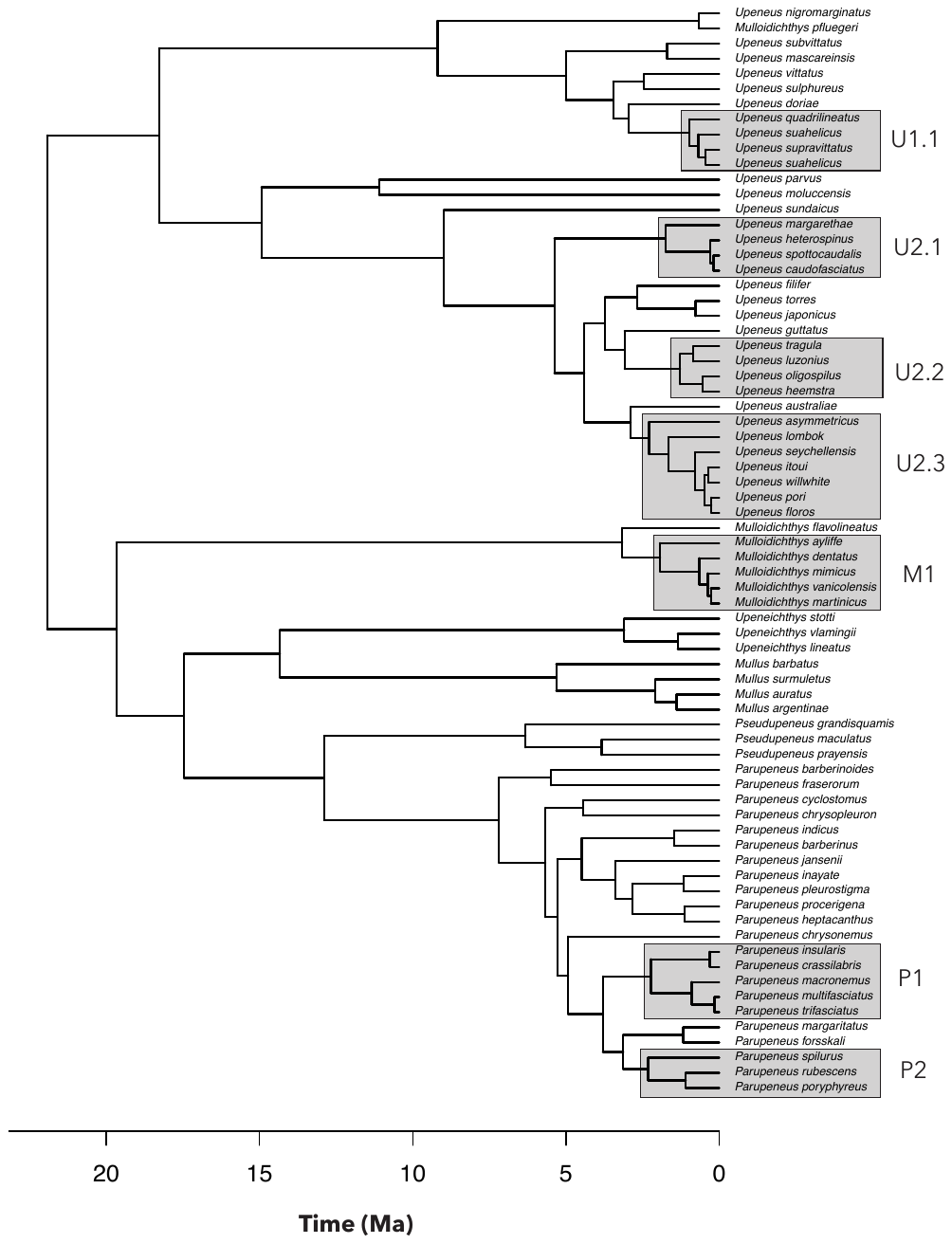

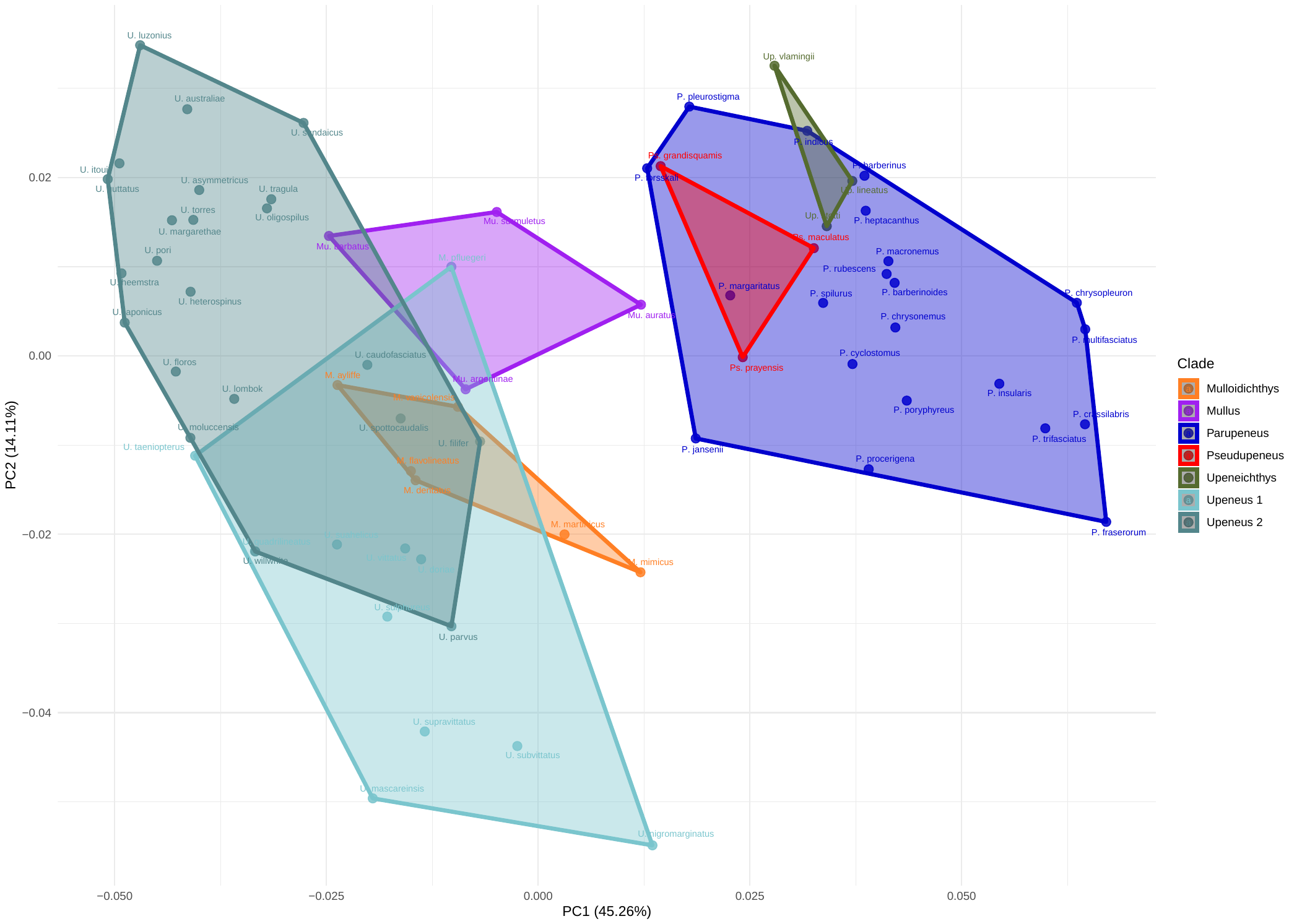

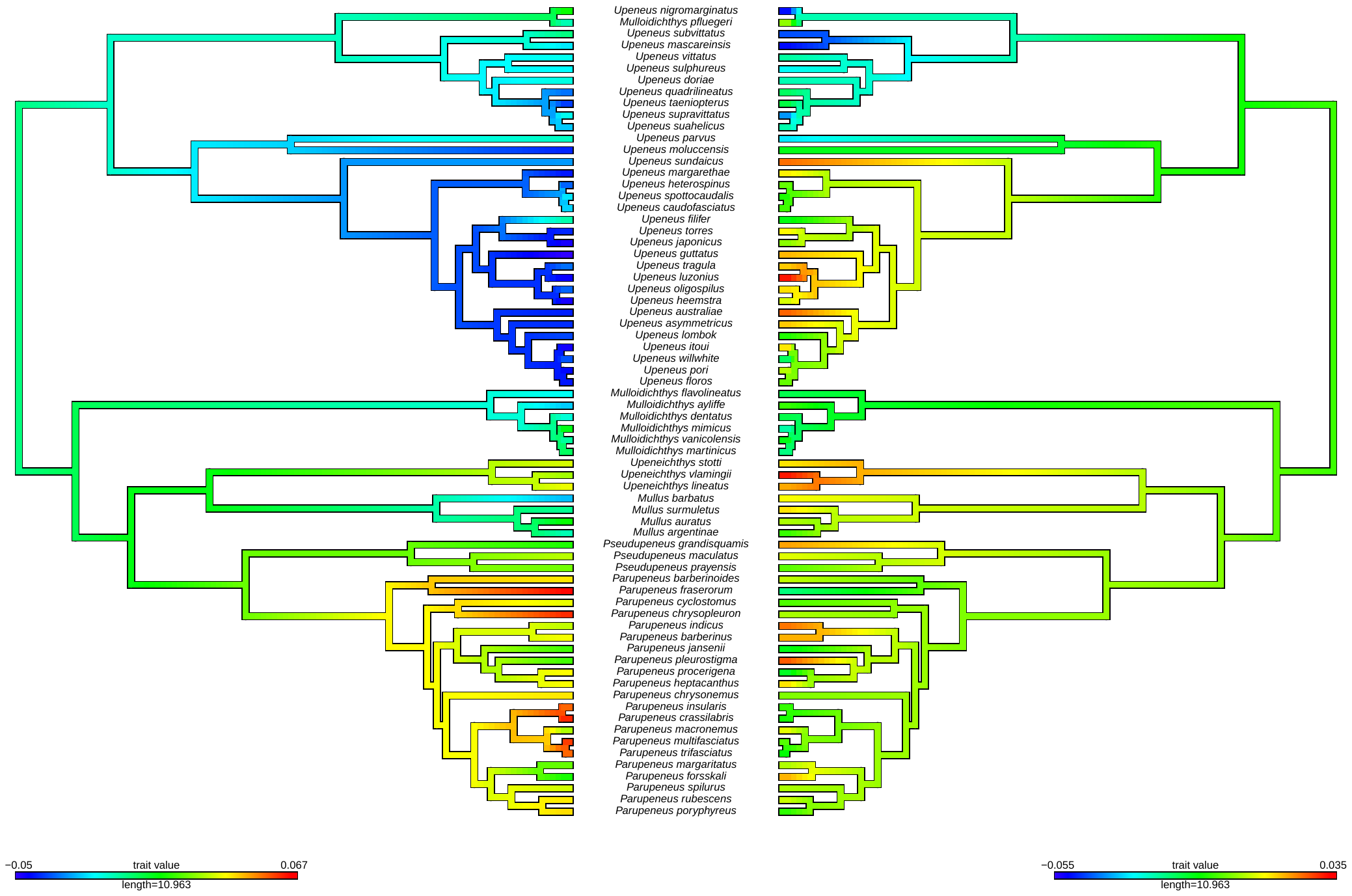

**Supplementary Fig. 2.** Phylogenetic placement of goatfish species groups. Location and labels of **s**pecies groups within the Mullidae, shown on the time-calibrated BEAST phylogeny. We define species group as a clade of three or more species whose mean node age is within the Pleistocene (2.58 - 0.012 Ma). Posterior probability values for this phylogeny can be found in Figure 3 and the HPD values are shown in Figure 2.

**Supplementary Fig. 3.** Body shape morphospace for goatfishes labeled by clade membership. Axis 1 and 2 from Principal Component Analysis (PCA) on Procrustes aligned body shape landmarks for 70 species. Each coordinate represents the mean shape of each species. PC1 represents 44.95% of the total variation, and PC2 represents 14.06% of the total variation. Species are grouped by clade, and this is indicated by color coded hulls to show the relative distribution of each clade in morphospace.

PC2

PC1

**Supplementary Fig. 4.** ContMap showing the distribution of PC1 and PC2 scores across the phylogeny. Higher values are indicated in warmer colors and lower values are shown in cooler colors. Figure generated using Phytools (Revell 2012).

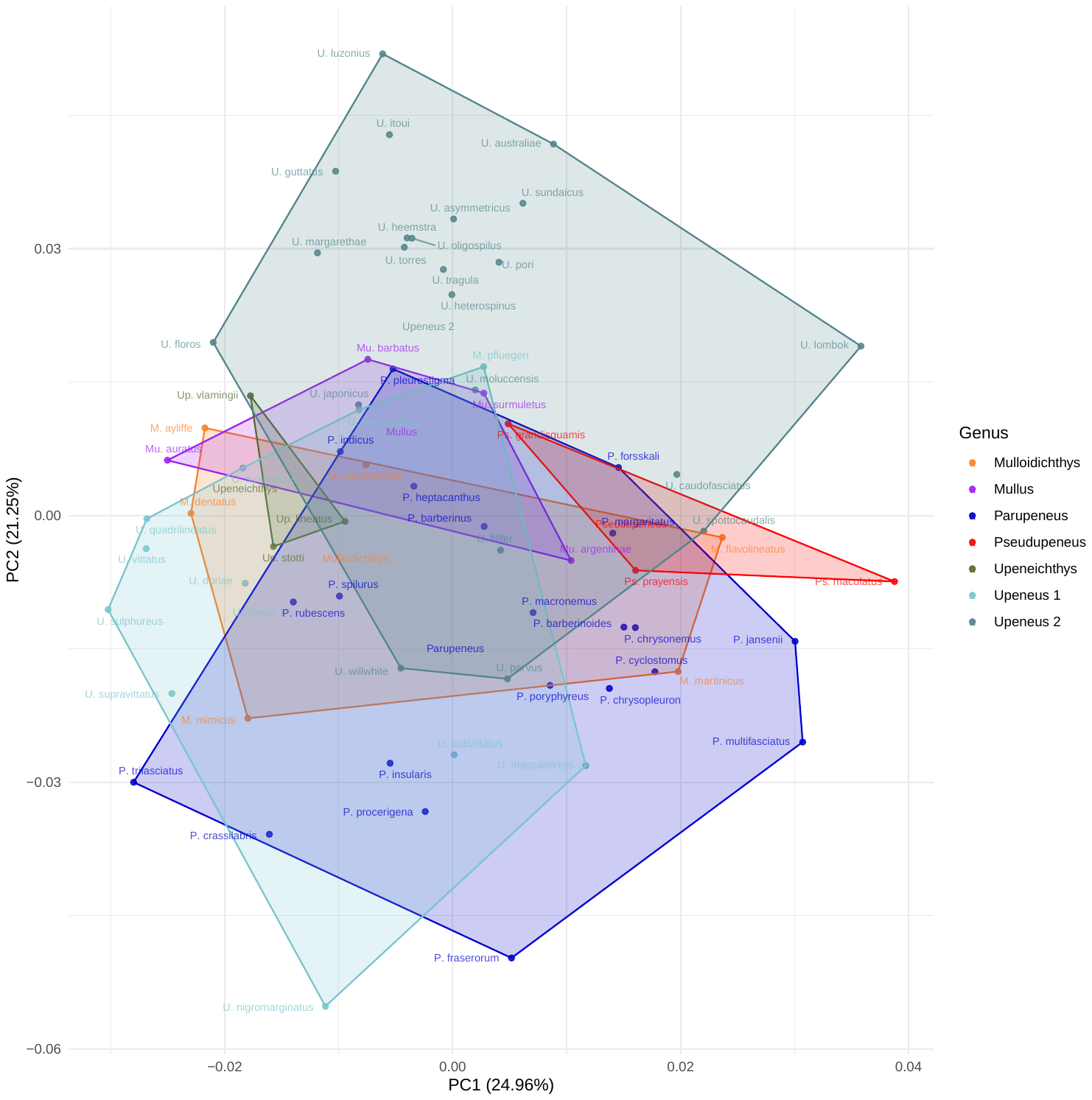

**Supplementary Fig. 5.** Phylogenetic PCA of goatfish body shape. Phylogenetic PCA using GLS-centering and projection implemented in the *gm.prcomp* function in geomorph 4.0.1 (Revell 2009; Adams and Otárola-Castillo 2013). Coordinates associated with each species and hulls are color coded by clade membership.

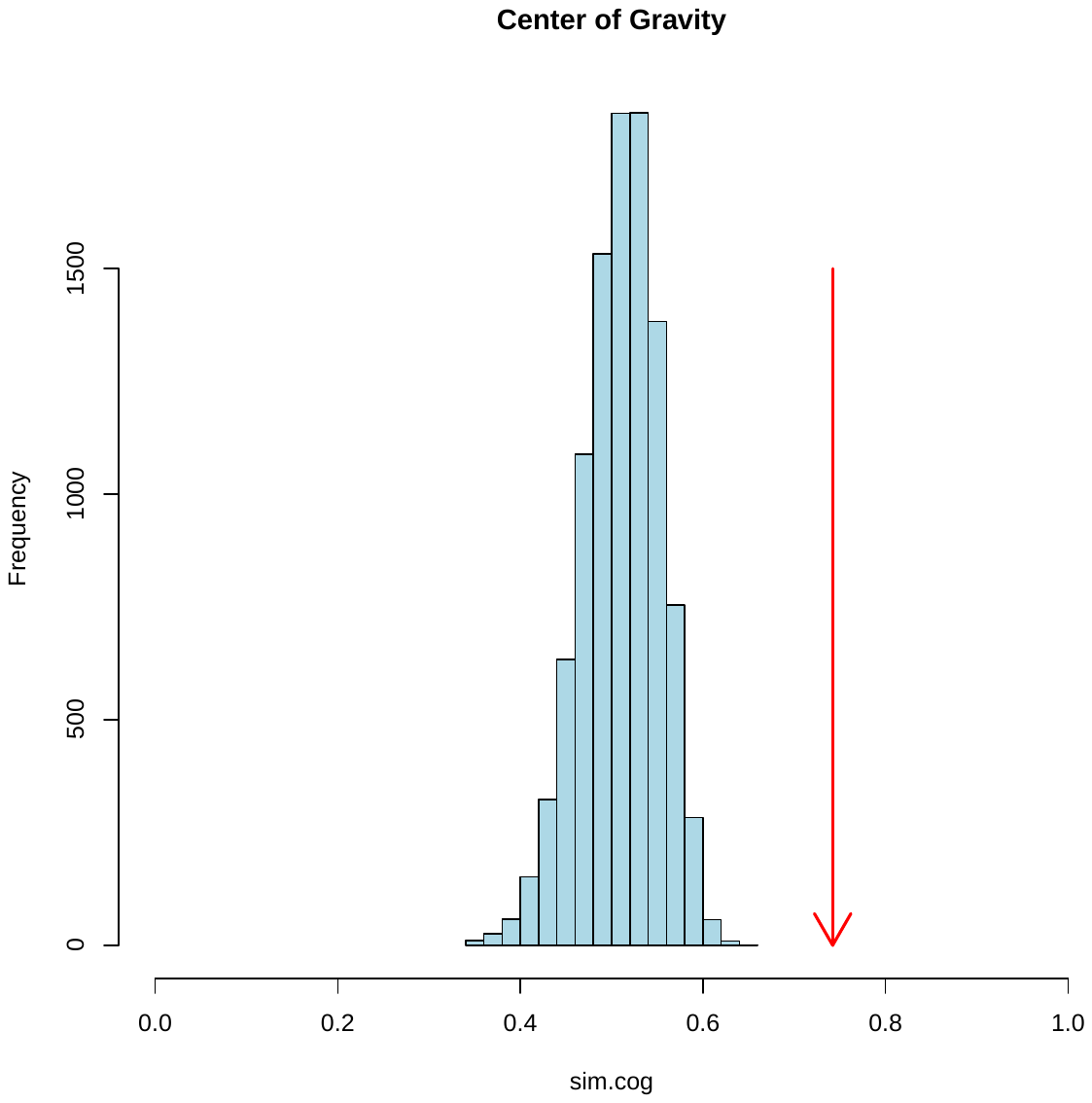

**Supplemental Fig. 6.** Weighted average times, termed Center of Gravity (CoG). The blue histogram are the results of 9,999 simulations generated under the assumptions of BM, and the red arrow indicates the CoG value for this study.

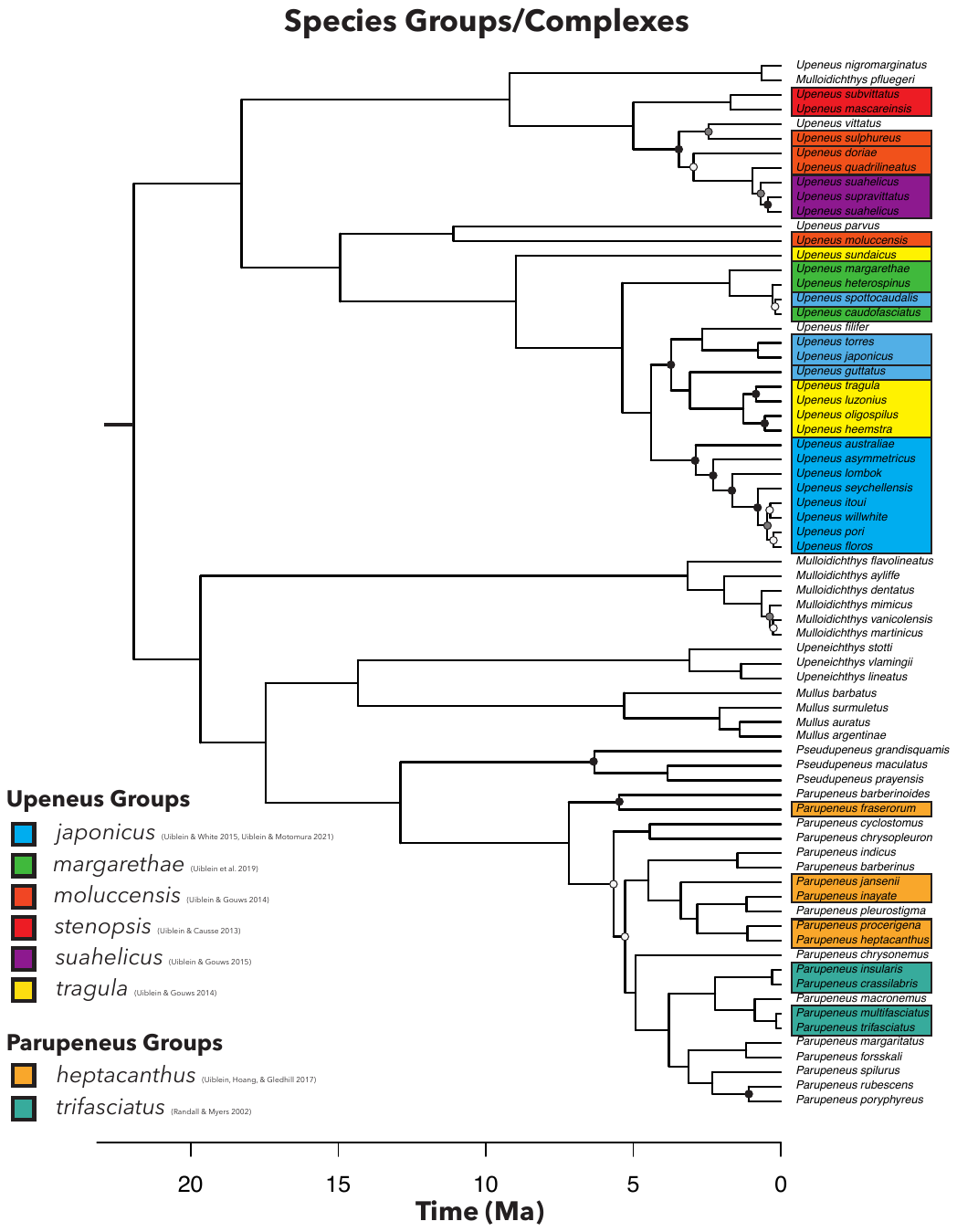

**Supplementary Fig. 7.** Described species complexes within the Mullidae. Distribution of previously described species groups and complexes on the time calibrated phylogeny inferred in this study. Species groups/complexes are indicated by color, and a black bar separating species within the same species group indicates that they are not monophyletic. Nodes without a circle have posterior probability of 1.0 from the BEAST analysis. Nodes with a circle are color coded by their posterior probability, with black = .99 - .95, grey = .94-.85, and white = <0.85.

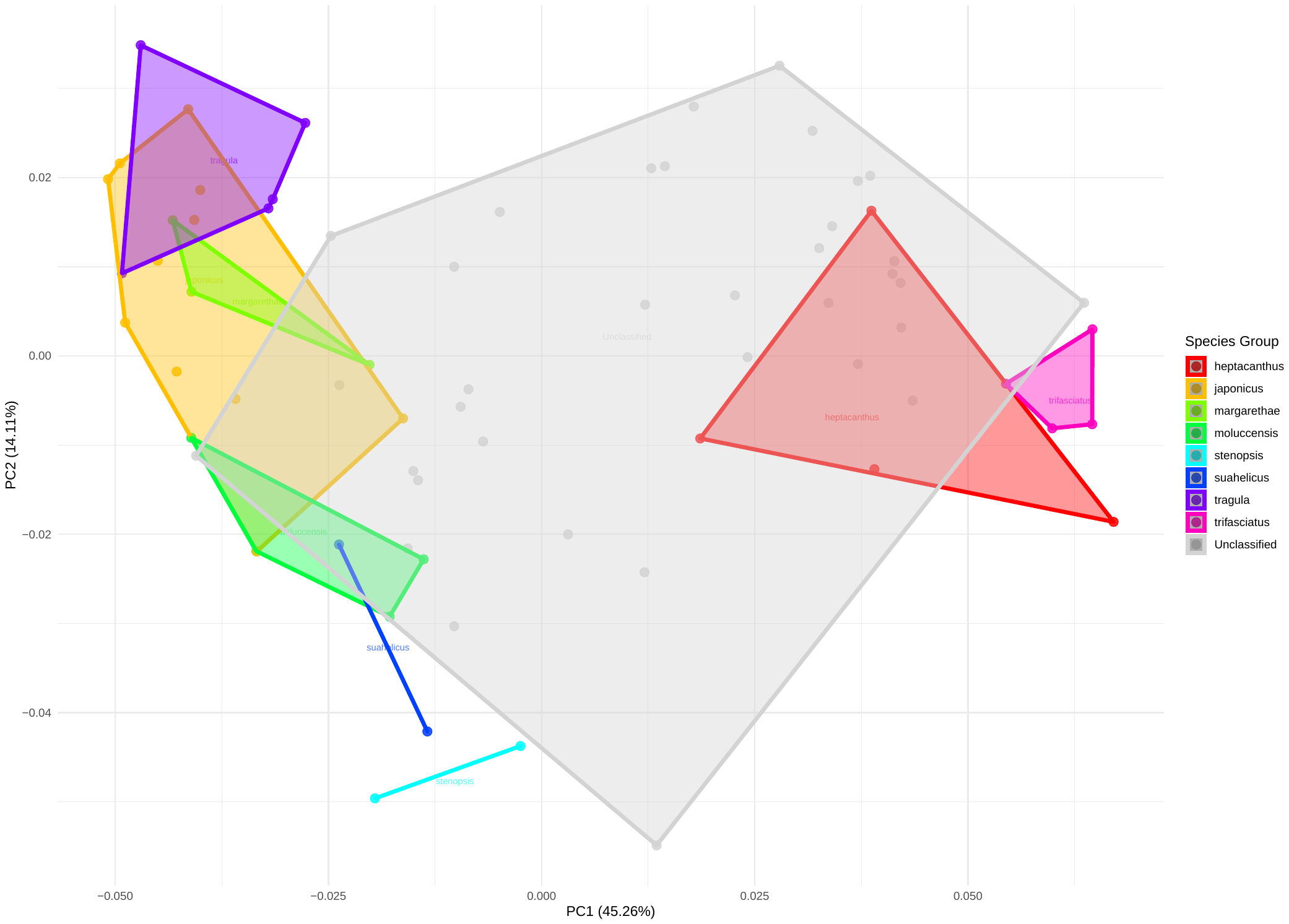

**Supplementary Fig. 8.** Body shape morphospace for the Mullidae labeled by species complex membership. Axis 1 and 2 from Principal Component Analysis (PCA) on Procrustes aligned body shape landmarks for 70 species. Each coordinate represents the mean shape of each species. PC1 represents 44.95% of the total variation, and PC2 represents 14.06% of the total variation. Species are grouped by membership in each species group/complex as defined by the literature (Fig. S7), and this is indicated by color coded hulls to show the distribution of group in morphospace. Species that are not assigned to a complex or group are indicated in light grey.
