## Appendix A Fossil Calibration Information for "Phylogenomics and body shape morphometrics reveal recent diversification in the goatfishes (Syngnatharia: Mullidae)"

**Appendix 1: Fossil Calibrations**

We used the calibration scheme described in Santaquiteria et al (2021) for both the outgroup and within Syngnatharia calibraions. The nodes associated with each calibration are indicated on the phylogeny shown in Figure 2. Calibrations #1 – #7 match those used in Santaquiteria et al (2021), while calibrations #8 and #9 are unique to this study. With the exception of calibration #1, all age priors were assigned a log normal distribution with the age of the fossil used as the hard lower bound. For the Syngnatharian root calibration (#1), we used a secondary calibration with a uniform distribution based on the protocol and description found in Santaquiteria et al (2021). Information about each calibration can be found below:

**Museum Collection Abbreviations**

FMNH: Field Museum of Natural History, Chicago, USA

IGM: Instituto Geológico de México, Mexico City, Mexico

PIN: Paleontological Institute, Moscow, Russia

MCSNV: Museo Civico di Storia Naturale di Verona, Verona, Italy

**1. Root Syngnatharia + Pelagiaria**

MRCA: *Taracichthys longipinnis* & *Parupeneus poryphyreus*

Secondary Calibration

103.5 Ma – 92 Ma (Santaquiteria et al. 2021)

**2. Syngnatharia**

MRCA: *Syngnathus louisianae* & *Eurypegasus draconis*

†*Gasterorhamphosus zuppichinii* MCSNV Na T 877

Syngnathiform according to (Near et al. 2012)

83.6 Ma, Campanian from Nardo, Italy (Santaquiteria et al. 2021)

Mean = 5.5, SD = 1.25

95% soft upper = 103

**3. Syngnathidae**

MRCA: *Syngnathus louisianae* & *Solenostomus cyanopterus*

†*Prosolenostomus lessinii* FMNH 94830

48.5 Ma, Porto Selvaggio, Lecce province, Italy (Sorbini 1981)

Mean = 9.8, SD = 1.25

95% soft upper = 83.6

**4. *Hippocampus***

MRCA: *Hippocampus abdominalis* & *Hippocampus kuda*

† *Hippocampus sarmaticus sp. nov.* T-245

11.6 Ma, Tunjice Hills, Slovenia (Žalohar et al. 2009)

Mean = 10.3, SD = 1.25

95% soft upper = 21.9

**5. Aulostomoidea (Aulostomidae)**

MRCA: *Aulostomus chinensi & Aeoliscus strigatus*

†*Eekaulostomus cuevasae* IGM 4716

61.5 Ma, Belisario Domínguez Quarry, Chiapas, México (Cantalice and Alvarado-Ortega 2016)

Mean = 6.2, SD = 1.25

95% soft upper = 83.7

**6. Fistularidae**

MRCA: *Fistularia corneta* & *Aulostomus chinensis*

†*Urosphen dubius* FMNH 94830

48.5 Ma, Pesciara site, Bolca Lagerstätte, northeastern Italy (Blainville 1818)

Mean = 3.7, SD = 1.25

95% soft upper = 61.7

**7. Pegasidae**

MRCA: *Pegasus volitans & Dactylopterus volitans*

†*Ramphosus rostrum* FMNH 94830

48.5 Ma, Pesciara site, Bolca Lagerstätte, northeastern Italy (Volta 1796)

Mean = 9.8, SD = 1.25

95% soft upper = 83.6

**8. Callionmymoidei (Callionymidae)**

MRCA: *Callionymus limiceps & Parupeneus poryphyreus*

*†Gilmourella minuta* MCSNV T.381/T.382

48.5 Ma, Pesciara site, Bolca Lagerstätte, northeastern Italy (Carnevale et al. 2019)

Mean = 9.8, SD = 1.25

95% soft upper = 83.6

**9. *Mullus***

MRCA: *Mullus argentinae & Upeneichthys stotti*

*†Mullus sp.* PIN 5073-10

13.0 Ma, Tsurevsky, North Caucus, Russia (Carnevale et al. 2006)

Described within the modern genus *Mullus* based on synapomorphies in Carnevale et al. (2006)

Mean = 10.0, SD = 1.25

95% soft upper = 48.8

Volta, G. S. 1796. Ittiolitologia Veronese del Museo Bozziano ora annesso a quello del Conte Giovambattista Gazola e di altri gabinetti di fossili veronesi. Verona.

Žalohar, J., T. Hitij, and M. Križnar. 2009. Two new species of seahorses (Syngnathidae, Hippocampus) from the Middle Miocene (Sarmatian) Coprolitic Horizon in Tunjice Hills, Slovenia: The oldest fossil record of seahorses. Ann. Paleontol. 95:71–96.
