## Appendix B Morphological Landmarks for "Phylogenomics and body shape morphometrics reveal recent diversification in the goatfishes (Syngnatharia: Mullidae)"

Appendix 2: Morphometric Landmark Scheme

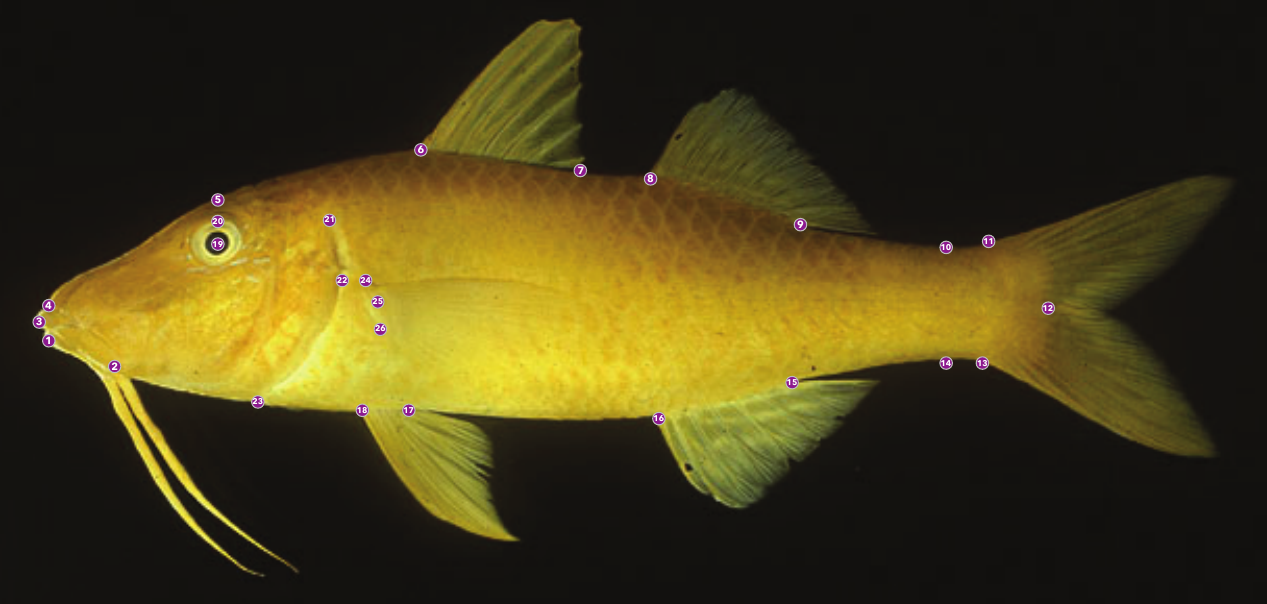

2. Tip of Lower Lip
3. Mandible Fork
4. Tip of Upper Lip
5. Snout Tip
6. Forehead Above Eye
7. Anterior First Dorsal Fin Base
8. Posterior First Dorsal Fin Base
9. Anterior Second Dorsal Fin Base
10. Posterior Second Dorsal Fin Base
11. Minimum Caudal Peduncle Width - Dorsal
12. Caudal Fin Base - Dorsal
13. Center Caudal Peduncle
14. Caudal Fin Base - Ventral
15. Minimum Caudal Peduncle Width – Ventral
16. Posterior Anal Fin Base
17. Anterior Anal Fin Base
18. Posterior Pelvic Fin Base
19. Anterior Pelvic Fin Base
20. Center of the Pupil
21. Top Center of the Eye
22. Posterior Postemporal
23. Posterior Opercule
24. Posteroventral Interopercal
25. Dorsal Pectoral Fin Base
26. Median Anterior Pectoral Fin Membrane
27. Ventral Pectoral Fin Base
